# Structure-guided design and development of cyclic peptide allosteric activators of Polycomb Repressive Complex 2

**DOI:** 10.1101/2025.06.26.661818

**Authors:** Yasuaki Tokodai, Mark Matyas, Azumi Ishigamori, Erin O’Connor, Angel Zhang, Sora Yamazaki, Tsunehide Hiraki, Noah Lohar, Liqi Yao, Ryan Stoner, Luis E. Paez-Beltran, Kayoko Kanamitsu, Noriko Nakaya, Satoshi Ichikawa, Justin Brumbaugh, Vignesh Kasinath, Fumika Yakushiji

## Abstract

Dysregulation of the histone methyltransferase Polycomb repressive complex 2 (PRC2) results in aberrant silencing of tumor suppressors and activation of oncogenes. Targeting PRC2 with compounds holds significant potential for both basic research and therapeutic applications. Here, we leveraged extensive structural studies of PRC2 to design a cyclic peptide that robustly activates PRC2. Structure-activity relationship studies guided the functional optimization of this cyclic peptide, yielding a Phenylalanine-type (Phe-type) cyclic peptide with approximately eight-fold activation compared to that of the poised state of PRC2. A 3.3Å cryo-electron microscopy structure of the PRC2–peptide complex, combined with biochemical analyses, revealed a shift in the H3K27 methylation from mono-(me1) and dimethylation (me2) to trimethylation (me3). Finally, we demonstrated that the cyclic peptide exhibits improved mouse plasma stability and can also be readily taken up by cells which results in a shift of the H3K27 methylation landscape to trimethylation, similar to the observed effects in vitro. These findings support the utility of such molecules for probing PRC2 activation and targeting dysregulated H3K27 methylation in cancer.

## Introduction

Histone post-translational modifications (hPTMs) are essential epigenetic modifications in cells and are closely associated with gene expression^1^. Site-specific hPTMs, established by dedicated enzyme complexes, define distinctive cellular states in response to environmental cues. Among these, the histone lysine (Lys)-specific methyltransferase Polycomb repressive complex 2 (PRC2) is well characterized for its role in gene silencing and stem cell regulation^2–7^. Consequently, its dysregulation leads to aberrant silencing of tumor suppressors and activation of oncogenes, highlighting its relevance as a cancer target^8–13^. Core PRC2 comprises EZH2, RBAp46/8, EED, and SUZ12^7,14–16^. The EZH2 subunit catalyzes sequential mono-, di-, and trimethylation of Lys 27 on the histone H3 *N*-terminal tail (H3K27) via its SET domain. Trimethylated H3K27 (H3K27me3) serves as a repressive chromatin mark critical for facultative heterochromatin formation. PRC2 also interacts with accessory cofactors in context-specific manners and exists in two mutually exclusive functional forms; PRC2.1 and PRC2.2^17,18^. PRC2.2 consists of two additional cofactor proteins JARID2 and AEBP2, which promote PRC2 recruitment to chromatin through ubiquitinated H2A Lys 119 (H2AK119ub) and allosterically activate EZH2 via trimethylated JARID2 (Jarid2K116me3). This modification establishes “Polycomb” regions of repressed chromatin^5,19–22^. JARID2K116me3 triggers EZH2 activation, driving H3K27 trimethylation and reinforcing a positive feedback loop that sustains Polycomb-mediated gene repression^23,24^.

Structural characterization of PRC2 bound to minimal stimulatory peptides, as well as to JARID2 and AEBP2, has revealed the mechanism by which H3K27me3 and JARID2K116me3 allosterically activate EZH2 methyltransferase activity^25–28^. Beyond this unique allosteric regulation, extensive structural insight into PRC2 have enabled the rational design of compounds to both activate and inhibit the complex. Herein, we addressed design, synthesis, and evaluation of cyclic peptide–based activators derived from the linear peptide motifs of endogenous activators such as H3K27me3 and JARID2K116me3, with the intent to improve their physicochemical properties, including flexibility, limited cellular permeability, and activation potential. Cyclic peptides with small ring sizes [< 10 amino acids (aa)], typically ranging from 500 to 1500 Da, exhibit greater resistance to proteolytic degradation^29,30^ and enhanced cellular permeability, with some achieving clinical success^31,32^. Hence, we first truncated the lead linear peptide to identify the minimal sequence required for PRC2 activation, followed by macrocyclization to generate effective cyclic variants. Through a combination of chemical synthesis, biochemical assays, and cryogenic electron microscopy (cryo-EM) analysis, we establish a robust strategy for developing PRC2 allosteric activators. Notably, our cyclic peptide more efficiently shifts the histone H3 methylation landscape towards H3K27me3 than the naturally allosterically activated PRC2 complex containing JARID2K116me3.

## Results

### Identifying the essential motif required for PRC2 activation

To define the minimal motif necessary for PRC2 allosteric activation, we prepared a 12-residue peptide which originates from the trimethylated H3 tail (Table 1, compound **1**) containing the key residues that stably bound and visible in the structures of activated PRC2 (**Figure 1A**)^25–27^. In addition to the 12-amino acids H3K27me3-containing peptide (compound **1**), we synthesized *C*-terminally truncated derivatives (compounds **2** and **3**; residues 23–30/31) and a linear peptide with a one-residue *N*-terminal shift (compound **4**; residues 22–29) via solid phase peptide synthesis (see Methods and Supplementary Information).

**Figure 1.**
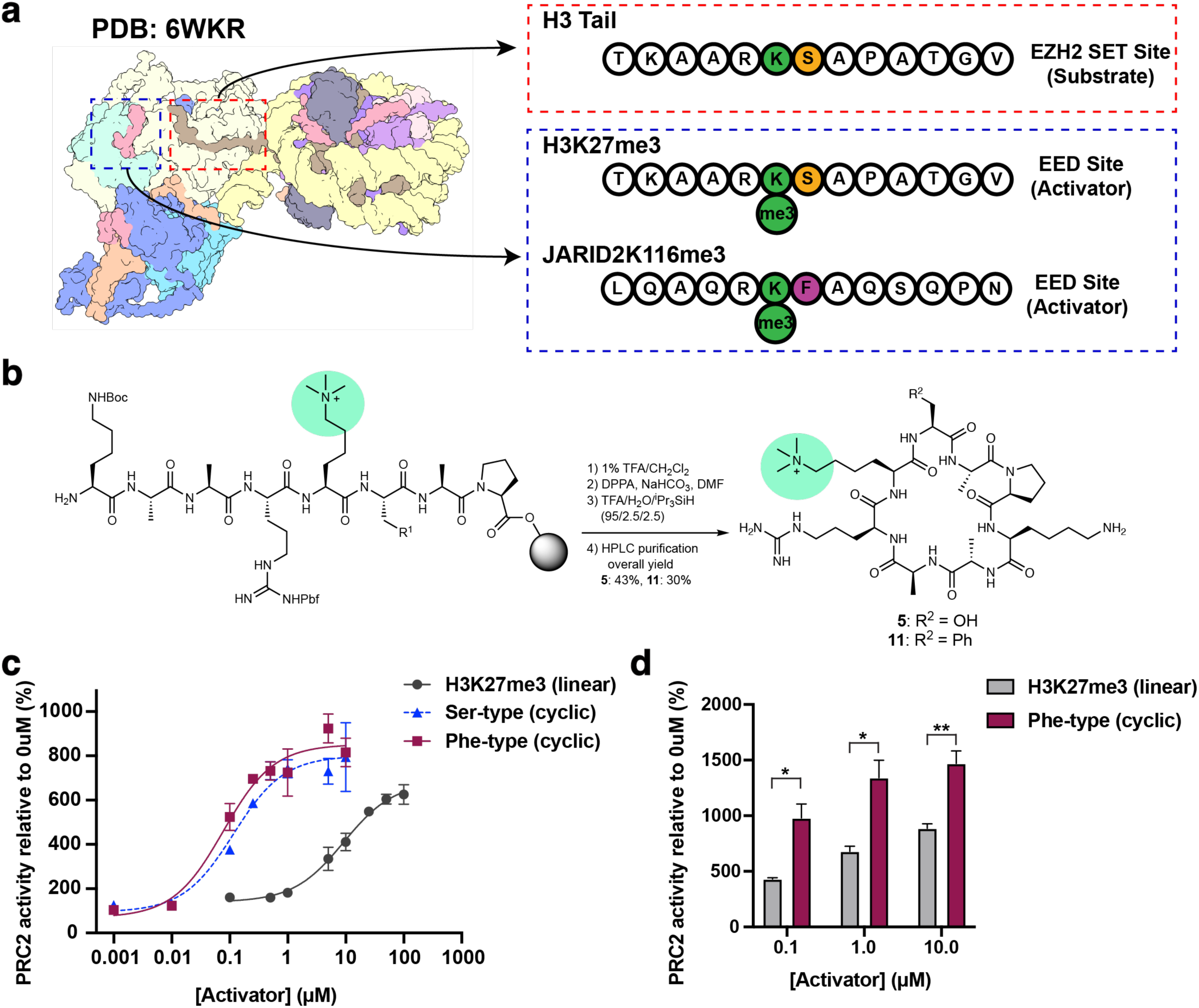
Design, synthesis and initial characterization of cyclic peptides targeting PRC2 allosteric activation. (**A**) Comparison of H3K27, H3K27me3, and Jarid2K116me3 peptide sequences that allosterically activate PRC2 from PDB: **6WKR**. (**B**) Outline of the chemical synthesis for the cyclo-derivatives. (**C)** Dose-response curves of either linear H3K27me3 (**1**) or Ser-type (**5**) or Phe-type (**11**) on PRC2 activity as measured by CPM counts. Histone H3 as substrate. n=3 technical replicates, error bars represent +/- s.e.m. (**D**) Increasing concentrations of activators (either compound **1** or **11**) and their effect on PRC2 activity. Human mononucleosomes as substrate. Welch’s two-tailed t-test, unpaired with equal variance assumed and no multiple testing correction. n=3 technical replicates. \**P* ≤ 0.05, \*\**P* ≤ 0.01

**Table 1.** Summary of PRC2 methyltransferase activity in the presence of different linear peptide activators. PRC2 activity (%) reported is the activity of PRC2 at 50 µM activator peptide. N=2 technical replicates were performed.

| comp. | peptide sequence | PRC2 activity (%) | EC<sub>50</sub> (μM)<br>[95% CI] |
| --- | --- | --- | --- |
| **1** | KAARK(me3)SAPATGG | 651 ± 29 | 9.23<br>[5.64 to 15.3] |
| **2** | KAARK(me3)SAPA | 583 ± 28 | – |
| **3** | KAARK(me3)SAP | 577 ± 1.9 | – |
| **4** | TKAARK(me3)SA | 485 ± 93 | – |

50 μM compound (n = 2)

The linear peptides (compounds **1**–**4)** were evaluated for *in vitro* activation of core PRC2 using histone H3 peptide (residues 21–44) as the substrate (Table 1). As expected, compound **1** stimulated PRC2 activity (651 ± 29% at 50 µM). The truncated peptides (compounds **2**–**4**) were tested to assess whether peripheral residue removal would impact PRC2 activation, aiming to reduce the overall molecular weight of a potential candidate. While all three peptides (compounds **2**–**4**) maintained the activity Compound **2**, lacking T-G-G from *C*-terminus of **1**, showed slightly reduced PRC2 activity (583 ± 28%). Further removal of *C*-terminal alanine (Ala; compound 3) had minimal impact (577 ± 1.9%). In contrast, *N*-terminal shifting (compound **4**) notably attenuated PRC2 activity (485 ± 93%). These results identified the KAARK(me3)SAP sequence in compound **3** as the minimal motif required for potent PRC2 allosteric activation.

### Macrocyclization and optimization of activating peptides

To enhance half maximal effective concentration (EC_50_) values, molecular stability, and cellular properties, we cyclized the essential PRC2 allosteric motif. A head-to-tail cyclized peptide based on the sequence of compound **3** was synthesized, yielding *cyclo*[KAARK(me3)SAP] [serine (Ser)-type activator; compound **5**] (**Figure 1B**). Cyclization was confirmed by mass spectrometry and high-performance liquid chromatography (HPLC) retention times (**Supplementary Figure 2**). Furthermore, nuclear magnetic resonance (NMR) hydrogen-deuterium exchange experiments indicated intramolecular hydrogen bonding at the Ala–Ala segment, stabilizing the peptide conformation, consistent with circular dichroism results. (**Supplementary Figure 3, 4**).

The activation of PRC2 by the Ser-type activator (compound **5)** was assessed *in vitro* using the same assay system described above. As shown in Table 2, the Ser-type activator exhibited enhanced PRC2 activator activity (730 ± 101% at 5 µM, EC_50_ = 0.116 µM) compared to that of the H3K27me3 linear peptide **1** (Table 1, EC_50_ = 9.23 µM). Cyclization of the linear peptide led to a marked improvement in PRC2 activation, with an approximately 80-fold improvement in EC_50_ (**Figure 1C**).

**Table 2.**
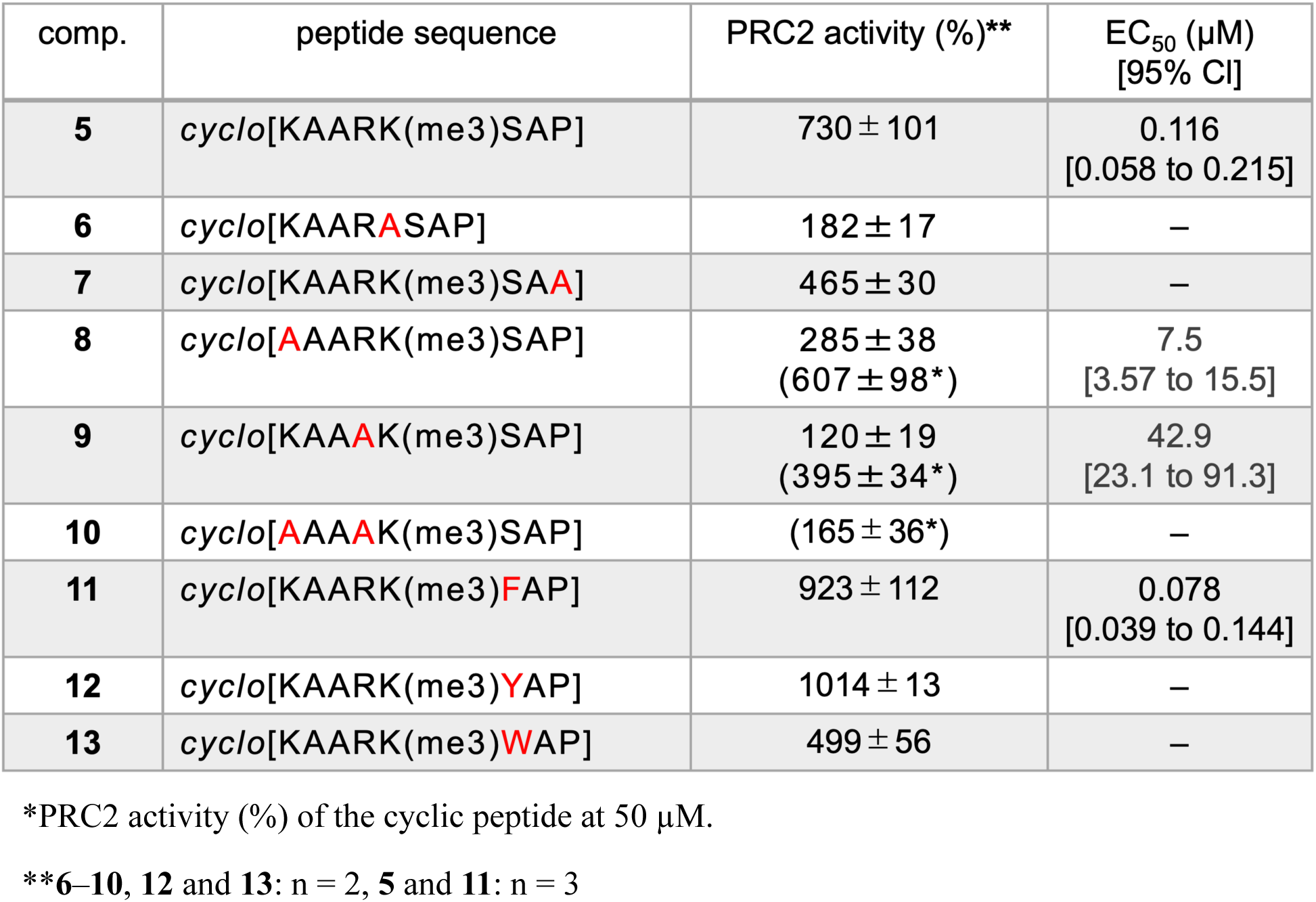
Summary of PRC2 methyltransferase activity in the presence of different cyclic peptide activators derived from compound **3**. PRC2 activity (%) reported is the activity of PRC2 at 5 µM cyclic peptide activator peptide. N=2 technical replicates were performed for compounds **6**-**10** and n=3 technical replicates were performed for compounds **5** and **11**.

To assess the contribution of individual residues to PRC2 activation, we performed Ala scanning on the Ser-type activator, substituting Lys(me3), proline (Pro), Lys or arginine (Arg) with Ala (compounds **6–9**). Substitution of Lys(me3), Lys, or Arg (compounds **6**, **8**, and **9**, respectively) significantly reduced activator activity (**Table 2**). Replacing both basic residues in compound **5** with Ala (compound **10)** resulted in a near-complete loss of activation potential. These results demonstrate that, in addition to Lys(me3), the peripheral Lys and Arg are critical for stimulating the PRC2 methyltransferase activity and likely contribute to key molecular interactions.

We next modified the Ser residue of compound **5** to aromatic residues to mimic the JARID2K116me3 sequence. Substitution with phenylalnine (Phe) yielded compound **11** (Phe-type activator), which enhanced PRC2 activation and significantly improved the EC_50_ (923 ± 112% at 5 µM, EC_50_ = 0.078 µM). Incorporating tyrosine (Tyr; compound **12**) also produced potent PRC2 activation, whereas tryptophan (Trp; compound **13**) reduced activity, likely due to steric hindrance or electrostatic incompatibility from the Trp side chain.

We then directly compared PRC2 activation by the Phe-type activator (compound **11**) and the linear H3K27me3 counterpart using recombinant mononucleosomes. As expected, the Phe-type showed superior activation across 0.1-10 µM concentrations (**Figure 1D**). Based on these findings, we selected the Ser- and Phe-type activators as lead candidates for further structural and biochemical analyses.

### Mechanism of allosteric activation of PRC2 by cyclic peptides

We first performed *in vitro* methyltransferase assays using Ser- and Phe-type activators with PRC2 containing JARID2 (aa 119-450) and AEBP2. Our previous structural studies have demonstrated that this six-subunit PRC2 complex (PRC2-AJ_119-450_) is more stable than core-PRC2 for cryo-EM structural analysis and remains in a poised state in the absence of an allosteric activation cue (e.g., JARID2K116me3 or H3K27me3)^26,33^ (**Figure 2A, B and Supplementary Figure 3A**). Both compounds strongly activated PRC2-AJ_119-450_ on purified histone H3.1 (aa 1-130) and mononucleosome substrates (**Figure 2C–F and Supplementary Figure 5**). While PRC2-AJ_119-450_ activity on H3 substrates was comparable to that of core PRC2, upon stimulation by the activator compounds, activation on nucleosomes was substantially higher (∼ 5 to 10-fold higher), consistent with nucleosomes being the physiologically relevant substrates *in vivo*^22^. Unlike histone peptides, nucleosomes bearing H3K27me3–once generated– by PRC2 activity act as potent allosteric activators and may compete with our cyclic peptides to increase the methylation levels. This competition likely explains why PRC2 activation by the cyclic peptides on nucleosomes is approximately 5–10-fold higher than that by histone peptide substrates.

**Figure 2.**
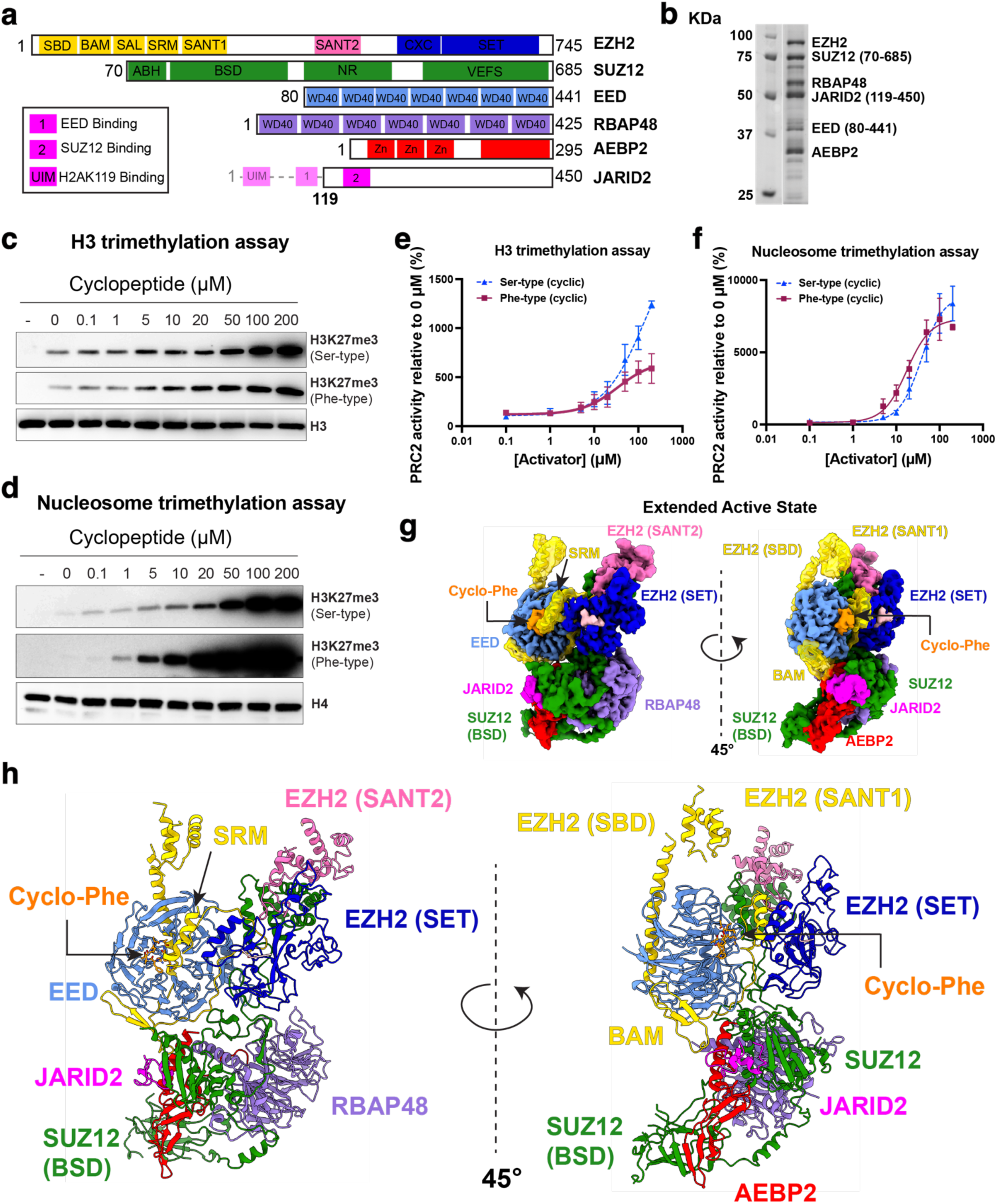
Cryo-EM structure of PRC2-AJ_119-450_ bound to Phe-type cyclic peptide activator. **(A)** Schematic representation of the domains of the 6-subunit PRC2-AJ_119-450_. **(B)** Representative Coomassie stained gel of purified PRC2-AJ_119-450_. **(C, D)** Representative immunoblots of H3K27me3 upon addition of cyclopeptide. Unmodified histone H3 or H4 immunoblots were used as either histone peptide or nucleosome loading controls. (**E, F**) Western blot-based quantification of the methyltransferase assays using PRC2-AJ_119-450_ and either Ser-type activator (compound 5) or Phe-type activator (compound 11) on histone H3 (aa 1-130) and mononucleosome substrates, respectively. n=3 technical replicates for each. Error bars represent +/- s.e.m. **(G)** Cryo-EM density of PRC2-AJ_119-450_ captured in its extended active state. The Phe-type cyclic peptide activator density is shown in orange. **(H)** Model of the full PRC2-AJ_119-450_ with the Phe-type cyclic peptide activator (orange) built into the cryo-EM density with EED (cornflower blue), EZH2 SET domain (medium blue), EZH2 SANT2 domain (hot pink), EZH2 SRM, EZH2 SANT1, EZH2 SBD, and EZH2 BAM (gold), JARID2 (magenta), SUZ12 (forest green), AEBP2 (red). EZH2 ‘QLKK’ segment (beige).

To determine whether PRC2-AJ_119-450_ activation by the Phe-type activator mirrors the allosteric mechanism of endogenous activators (H3K27me3 or JARID K116me3), we resolved a 3.3 Å cryo-EM structure of PRC2-AJ_119–450_ bound to the Phe-type activator in two distinct active states – both with the cyclic peptide bound but distinguished by the presence or absence of an ordered SRM, reflecting the conformational flexibility of the SRM (**Figure 2G, H, Supplementary Figure 6, 7A-E**). Our earlier studies similarly revealed two active states of PRC2 based on SRM ordering^26,27^. In addition, we previously also reported PRC2-AJ_119–450_ bound to an inhibitory RNA lacking an ordered stimulatory response motif (SRM)^33^. Notably, in the SRM-ordered state of our structure, we observed a previously unassigned density in the EZH2 (SET) catalytic domain, likely representing a segment of EZH2 undergoing automethylation (**Supplementary Figure 7F**). Prior evidence suggests the QLKK segment (EZH2 aa 507-510) undergoes automethylation and enhances PRC2 methyltransferase activity^34,35^. As the cryo-EM complex contained no other substrates bearing H3K27 or JARID2K116, we assign this density to the QLKK segment within EZH2. Model building into the catalytic site supports this assignment, consistent with QLKK occupying the active site (**Figure 2G, H, and Supplementary Figure 7F, G**). In summary, we identify multiple active conformations of PRC2-AJ_119–450_, all with the cyclic peptide bound to EED, but differing in SRM ordering and presence of the automethylated EZH2 segment (aa 507–510).

We successfully built the cyclic peptide and most of PRC2 into our 3.3 Å cryo-EM reconstruction, revealing the molecular interactions between PRC2 and the Phe-type cyclic activator. The cyclopeptide is positioned within the EED pocket (buried surface area = 983 Å^2^), where its basic residues, Lys and Arg, form electrostatic interactions with acidic residues of the SRM (**Figure 3A**)^36^. These salt bridges would enhance stability of the interaction between the cyclic peptide and EZH2 (SRM). The Phe-type activator binds a similar EED site as linear activators (H3K27me3 or JARID2 K116me3; buried surface area = 1136 Å^2^), with its trimethylated Lys situated in an aromatic cage formed by EED F97, Y148, Y365, and W364 (**Figure 3B**). Furthermore, we observed a putative CH/π interaction between the Phe and EED’s P95, which may contribute to the enhanced activation seen with cyclic compounds **11** and **12** compared to that of compound **5**. Relative to the linear JARID2K116me3 peptide in prior structures, the geometry of the cyclic peptide directs nonmethylated Lys (K1) and Arg (R4) toward the SRM acidic residues (**Figure 3B and Supplementary Figure 8A, C**).

**Figure 3.**
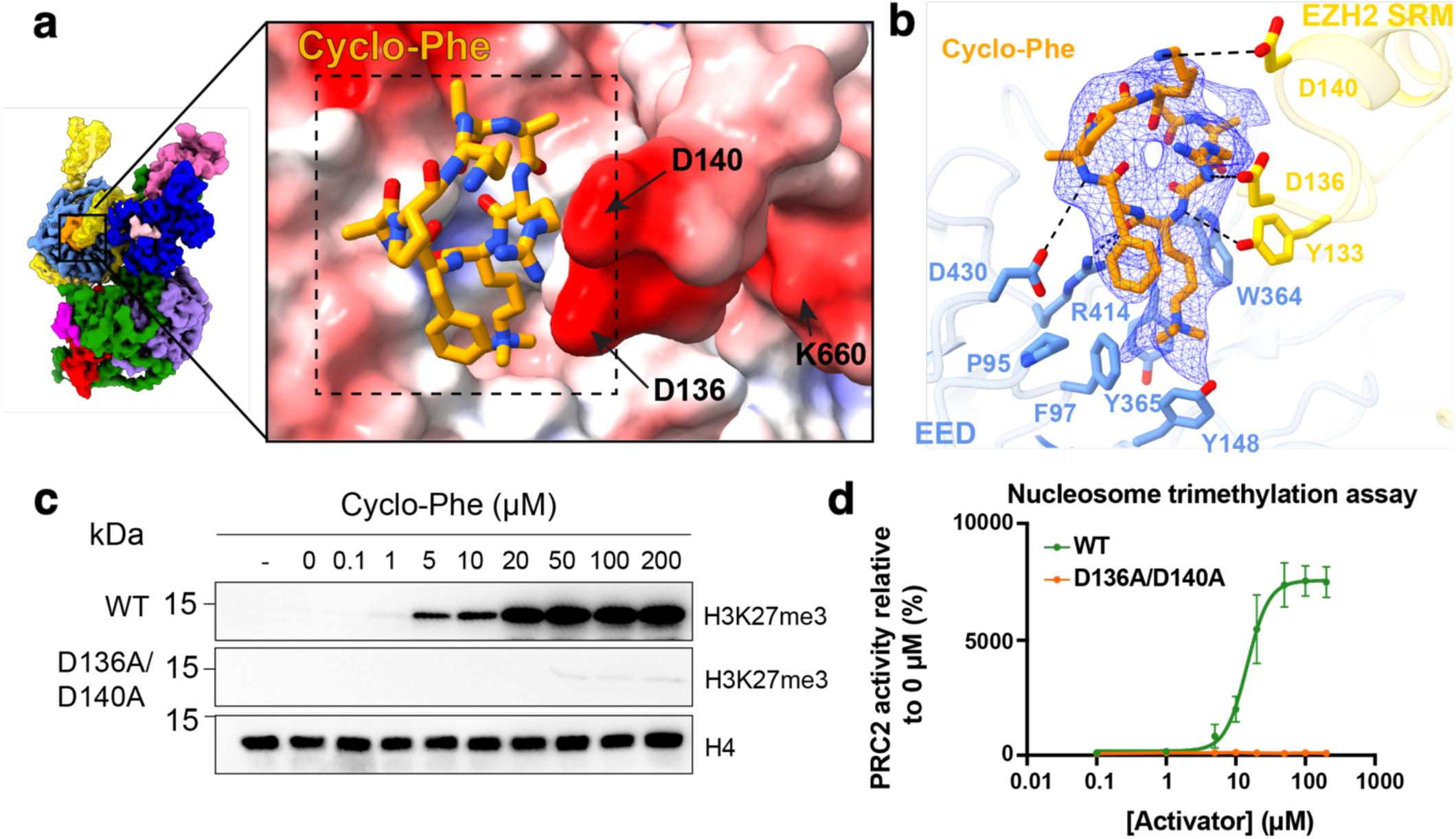
Interaction of Phe-type cyclic peptide activator with EED and EZH2 SRM. (**A**) Close-up view of Phe-type activator bound to PRC2-AJ_119-450_, showing electrostatic surface potential of EED pocket and D136, D140 of SRM and K660 of the SET helix. **(B)** cryo-EM density of Phe-type activator (shown as mesh) with notable side chain interactions of EED (R414, P95, F97, Y148, W364, and Y365) and EZH2 SRM (Y133, D136, D140). **(C, D)** Methyltransferase assays and quantification probing for H3K27me3 for both PRC2-AJ_119-450_ WT and PRC2-AJ_119-450_ D136A/D140A double mutant on mononucleosome substrates. Histone H4 is used as a nucleosome loading control. n= 3 technical replicates. Error bars represent +/- s.e.m.

To validate our structural findings and assess whether activation occurs via EZH2’s SRM, we performed methyltransferase assays using PRC2-AJ_119–450_ harboring point mutations (D136A and D140A) in the SRM that disrupt interaction with the Phe-type activator (**Supplementary Figure 8B**). Given that the Lys and Arg of the cyclopeptide extend toward D136 and D140 of EZH2, we hypothesized that mutating these aspartic acids (D136 and D140) to Ala would disrupt the critical cyclic peptide–EZH2(SRM)–EZH2(SET) interaction axis required for PRC2 activation. As expected, PRC2-AJ_119-450_ D136A/D140A showed nearly complete loss of activation by the cyclic peptide, with only minimal activity observed at Phe-type activator concentrations above 50 µM (**Figure 3C, D** and **Supplementary Figure 8D**). These biochemical results support a mechanism in which the cyclic peptide activates PRC2 through the SRM, similar to H3K27me3 and JARID2K116me3^26,27^.

### Methylation landscape of an allosterically activated PRC2

Mono-, di- and trimethylation of Lys 27 on histone H3 (H3K27) each play distinct roles in transcriptional regulation across the mammalian genome^37^. To assess the impact of the cyclic-Phe peptide on PRC2 activity, we employed an *in vitro* assay detecting H3K27me1, H3K27me2, and H3K27me3. Previous studies using a five-subunit PRC2 complex (EZH2, EED, SUZ12, AEBP2, RBAp48) demonstrated that, in the absence of stimulatory cofactors, PRC2 efficiently methylates H3K27me0 and H3K27me1, but modifies H3K27me2 with lower efficiency^37,38^. Consistent with these findings, our six-subunit PRC2 (PRC2-AJ_119–450_) showed the highest activity for catalyzing H3K27me1, with reduced efficiency for H3K27me2, using unmodified nucleosomes as substrate (**Figure 4A**). As a positive control, we tested PRC2-AJ_1-450_ construct–– a six-subunit PRC2 complex containing JARID2(1-450) ––which is intrinsically allosterically activated. This complex efficiently deposited mono- and dimethylation marks on H3K27, with modest levels of H3K27me3, as confirmed by western blotting. Notably, addition of cyclic-Phe to PRC2-AJ_119–450_ enabled robust catalysis of both H3K27me2 and H3K27me3 from H3K27me1 (**Figure 4A, B**). This indicates that exposure to cyclic peptides enhances PRC’s ability to promote EZH2-mediated trimethylation of H3K27. However, structural comparison of the EZH2 SET domain in previously resolved PRC2 structures with the cyclic-Phe-bound form revealed no significant alterations in the Lys channel or the SET domain (r.m.s.d. = 0.78 Å) (**Supplementary Figure 9**). Thus, the cyclic peptide likely functions by stabilizing the SRM and SET helix or by maintaining PRC2 in a constitutively active confirmation, thereby favoring H3K27 trimethylation.

**Figure 4.**
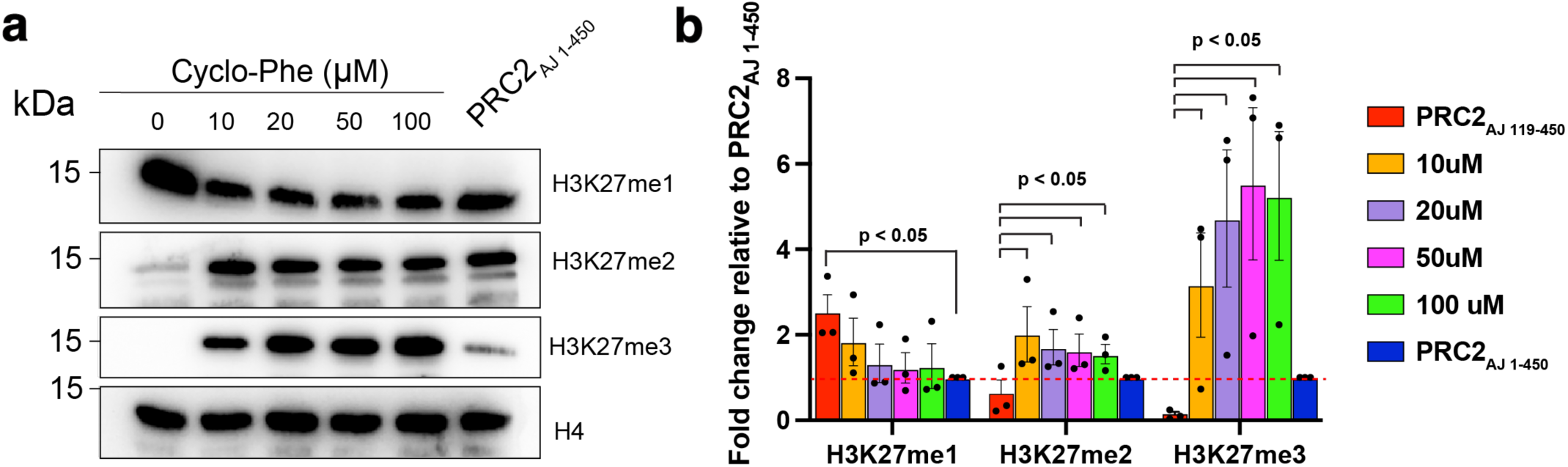
Comparison of the H3K27 mono-, di-, and tri-methylation landscape of PRC2-AJ_119-450_ with Phe-type cyclic peptide activator and PRC2-AJ_1-450_. **(A)** Western blot methyltransferase assays of H3K27me1, H3K27me2 and H3K27me3 for increasing amounts of Phe-type cyclic peptide and the intrinsically allosterically active PRC2-AJ_1-450_. n=3 technical replicates. Histone H4 is used as a nucleosome loading control. **(B)** Quantification of western blots, normalized to PRC2-AJ_1-450_. H3K27me1 is statistically significant between PRC2-AJ_119-450_ and PRC2-AJ_1-450_. Addition of cyclic peptide results in statistically significant difference in H3K27me2 and H3K27me3 signal when tested to PRC2-AJ_119-450_. Welch’s two-tailed t-test, unpaired with equal variance assumed and no multiple testing correction. n=3 technical replicates.

### Mouse plasma stability and cellular uptake of Phe-type cyclic peptide activator

The therapeutic potential of our cyclic peptides depends on favorable *in vivo* properties, such as cellular uptake and plasma stability. To evaluate this, we performed peptide-based transfections of compound **11** in suspension cultured HEK293 cells and assessed mouse plasma stability for both a linear H3K27me3 peptide (compound **3**) and our cyclic derivatives (compounds **5** and **11**). To assess the suitability of our cyclic peptides for use in animal tissues, we evaluated their stability in mouse plasma. The Ser-type activator (compound **5**), Phe-type activator (compound **11**), and a linear H3K27me3 peptide (compound **3**) were incubated in mouse plasma. Mass spectrometry analysis revealed that both cyclic activators exhibited ∼30% greater resistance to proteolysis than the linear peptide did (**Figure 5A and Supplementary Figure 10, 11**). These results indicate that our macrocyclization strategy effectively enhances peptide stability for higher-order biological applications.

**Figure 5.**
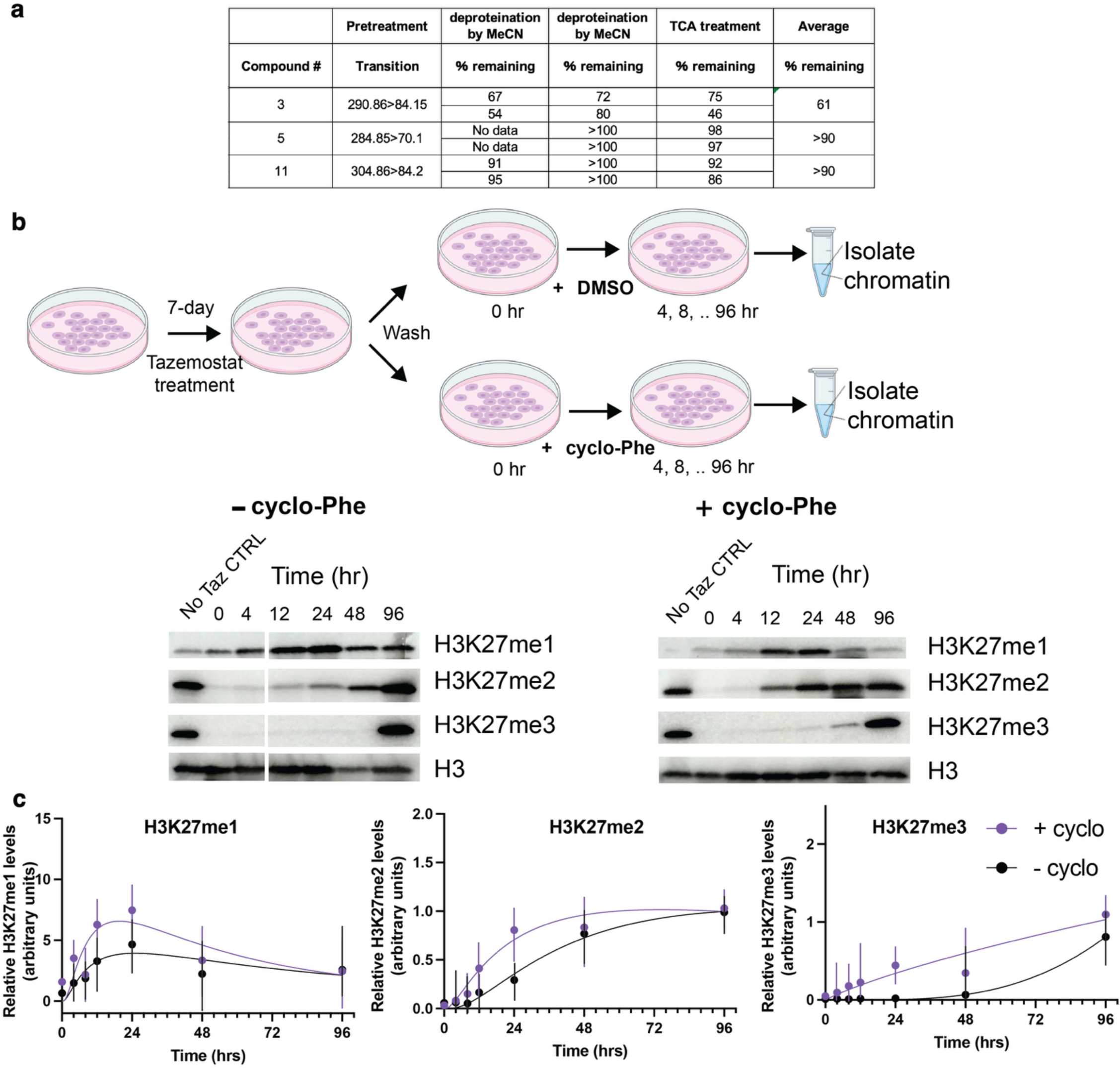
Cellular uptake and stability of Phe-type activator. **(A)** Summary of mouse plasma stability assay between linear H3K27me3 (compound **3**) and cyclic peptides (**5** and **11**). Conditions for data collection are listed in the methods. N = 4–6. Chemical structure of the FITC-labeled cyclic peptide used for cellular studies. The non-methylated lysine was replaced with an appended FITC molecule. (**B**) (top) Experimental schematic of the transfection-based assay in mESCs to gauge the recovery of H3K27me1/2/3 after Tazemostat treatment. (bottom) Representative western blot comparing the H3K27me1/2/3 in mESCs treated with DMSO or a single dose of cyclo-Phe activator. N=4 replicates were performed. (**C**) Quantification of the H3K27me1/2/3 recovery profiles in mESCs with or without cyclo-Phe treatment from all four replicates.

Next, to investigate the cellular uptake of the cyclic peptide and its effect on H3K27me3 levels in cells, we first synthesized a fluorescein isothiocyanate (FITC)-labeled Phe-type activator by modifying the nonmethylated Lys (**Supplementary Figure 12A**). Suspension HEK293 cells were either left untreated (DMSO) or transfected with the FITC-labeled peptide using a peptide-based transfection reagent (50 µM in dimethyl sulfoxide [DMSO]; see Methods). After one cell cycle (∼24-48hrs), both transfected and control cells were subjected to flow cytometry to quantify FITC-labeled peptide uptake (**Supplementary Figure 12B**). Cells were gated for single, live populations, followed by gating along the 488 nm axis to identify FITC+ cells. Nearly all transfected cells showed a shift along the FITC x-axis, indicating uptake. Applying stringent gating criteria, we conservatively estimated that at least 23% of transfected HEK cells internalized the cyclic peptide (**Supplementary Figure 12C**). Separately, we imaged HT-29 cells transfected with the FITC-labeled cyclic peptide in the presence or absence of the transfection reagent. Cell morphology remained unchanged, and most reagent treated cells were FITC+ (**Supplementary Figure 13**).

To investigate the effect of cyclic peptide uptake on H3K27me1/2/3 levels in cells, we first examined the levels of H3K27me1/2/3 in both HEK293 and mouse embryonic stem cells (mESCs). However, both cell lines already exhibited high levels of H3K27 methylation and could therefore not be used directly to examine the effect of the cyclic peptide on H3K27 methylation (**Supplementary Figure 12D, E**). Instead, we first completely depleted H3K27me1/2/3 in mESCs using tazemostat, followed by either transfection with a single-dose of Phe-type cyclic peptide activator or a DMSO control to examine the recovery and landscape of H3K27me1/2/3. We found that the Phe-type cyclic peptide activator facilitated faster recovery of H3K27 methylation. Moreover, H3K27me3 levels recovered more quickly and were higher than those in the DMSO control, mirroring our in vitro observations (**Figure 5B, Supplementary Figure 14**). These findings together support the feasibility of future cellular or *in vivo* studies–– particularly in disease-relevant models–– to examine how cyclic peptide uptake influences H3K27me3 levels and gene expression.

## Discussion

A substantial number of studies have explored small-molecule effectors of PRC2 as tractable targets for either loss -or gain-of-function mutations, which disrupt H3K27me3 levels and contribute to oncogenesis. Most efforts have focused on PRC2 inhibitors, characterized into five main classes: (a) EZH2 catalytic inhibitors (e.g., S-adenosyl-L-methionine (SAM) competitors^38,39^), (b) EED-inhibitors (disrupting EED-EZH2 interactions^40–42^), (c) EZH2 covalent inhibitors^43^, (d) EZH2 degraders^44^, and (e) EED degraders^45^. In contrast, few studies have pursued specific PRC2 activators, largely due to the complex allosteric activation mechanism of PRC2. Existing approaches primarily rely on histone H3 or JARID2 peptide mimetics^46^. Nonetheless, small molecules modulating PRC2 enzymatic activity have proven instrumental in dissecting its cellular roles. Thus, identifying a robust PRC2 activator, particularly in disease contexts where PRC2 activation represents a therapeutic vulnerability^47^, remains a worthwhile goal. Over the past decade, cyclic peptides have emerged as promising drug-like molecules, with increasing success in both academic and clinical settings^48^. However, to our knowledge, no cyclic peptides targeting PRC2 have been developed.

Here, we utilized structural insight into PRC2 to rationally design cyclic peptide allosteric activators. Using cryo-EM, we elucidated the molecular mechanism of activation by resolving the PRC2–activator complex. Structure-function analyses reinforce prior findings that allosteric activation involves coordinated interactions among EED, EZH2(SRM), and EZH2(SET), while offering more detailed mechanistic insight into how the cyclic peptide enhances PRC2-mediated histone methylation. Preliminary assessments of mouse plasma stability and cell permeability further support the cyclic peptide’s potential as a valuable chemical probe and a lead for medicinal chemistry targeting PRC2 dysregulation in cancers such as gliomas (H3 K-to-M mutations) and lymphomas^47^.

Loss of function mutations in core PRC2 components––EZH2, EED, and SUZ12––as well as accessory subunits such as JARID2 and AEBP2, are prevalent in various myeloid malignancies, positioning PRC2 as a critical target for anticancer therapy^49^. However, no PRC2 activators are currently available for laboratory or medicinal chemistry applications. Histone peptide–based activators offer a promising approach due to PRC2’s specific allosteric activation mechanism. Indeed, linear peptide activators have been developed for PRC2 harboring a loss-of-function EED(I363M) mutation^46^. However, their poor cell permeability necessitates further optimization to enable effective intracellular PRC2 activation.

Conversely, gain-of-function mutations in the catalytic SET domain of EZH2–– particularly Y641X (X=S, N, F, C, or H)––have been identified in lymphomas dependent on oncogenic EZH2 activity^49,50^, making PRC2 inhibition a viable therapeutic strategy. EZH2 inhibitors, including SAM-competitive agents like tazemetostat, have reached clinical trials and FDA approval for follicular lymphoma and epithelioid sarcoma. However, these inhibitors can induce *de novo* EZH2 mutations in lymphoma models, leading to resistance^51,52^. Despite the emergence of new PRC2 inhibitor classes, a deeper genetic understanding of PRC2 gain-of-function effects is essential for improved therapeutic strategies. Currently, there are no effective tools to allosterically activate PRC2 in a controlled manner within lymphoma models. The cyclic peptide developed in this study represents a promising candidate for probing PRC2 activation in such cellular contexts.

Clinical application of cyclic peptide activators will require enhanced cell permeability, potentially through side-chain modifications that do not disrupt EZH2(SRM) binding, and improved plasma stability via non-native peptide linkages to minimize degradation. Furthermore, additional studies are needed to investigate the impact of cellular uptake of these peptides on both global and locus-specific H3K27me3 levels, as well as associated changes in gene expression. Encouragingly, our lead candidates robustly activate PRC2 while retaining favorable *in vivo* properties.

Our approach, combining peptide-based compound development with cryo-EM structural analysis, has enabled the first successful creation of a cyclic peptide allosteric activator of PRC2. The mechanistic insights into PRC2 activation presented here offer a foundation for further modular optimization of activator compounds. Moreover, future studies using the Phe-type cyclic-peptide developed in this study may uncover new biological insights into malignancies driven by PRC2 gain-of-function mutations.

## Experimental Methods

### Expression and purification of PRC2 constructs

Full-length EZH2 isoform 2, EED, SUZ12, RBAp48, Strep-GFP-tagged embryonic isoform of AEBP2, and Strep-GFP-tagged truncated JARID2 (amino acids 119-450 or amino acids 1-450) were assembled into a single multi-bac plasmid using ligation independent cloning (438 series, Scott Gradia). The synthesized multi-bac plasmid was verified using full plasmid sequencing (Plasmidsaurus). Point mutations were generated by Genscript Corp. Baculovirus were made from the multi-bac plasmid using Sf9 and subsequently used for expression of PRC2 complex in Trichoplusia ni (HighFive) system. HighFive cells were transfected with baculovirus at 28°C for 66 hours, washed with cold PBS buffer, and frozen in liquid nitrogen until use. All purification steps were performed in a 4°C cold room. Cells were lysed in lysis buffer (25 mM HEPES pH 7.9 at 4°C, 250 mM NaCl, 2 mM MgCl_2_, 1 mM TCEP, 0.5% NP-40, 10% glycerol, protease inhibitor cocktail supplemented with aprotinin, leupeptin, and PMSF) for 1 hour and sonicated three times at 60% power with 10 sec ON and 20 sec OFF (Q-Sonica). Cell debris was then removed by centrifugation at 15,000 rpm for 35 minutes. The supernatant was incubated with Streptactin XT (IBA LifeSciences) overnight at 4C. The resin was washed with 10 column volumes (CV) of lysis buffer, 10 CV of high-salt wash buffer (25 mM HEPES pH 7.9 at 4°C, 1 M NaCl, 2 mM MgCl_2_, 1 mM TCEP, 0.01% NP-40, and 10% glycerol), and 20 CV of low-salt wash buffer (25 mM HEPES pH 7.9 at 4°C, 150 mM NaCl, 2 mM MgCl_2_, 1 mM TCEP, and 10% glycerol). Proteins were then eluted in elution buffer (25 mM HEPES pH 7.9 at 4°C, 150 mM NaCl, 2 mM MgCl_2_, 1 mM TCEP, 1X BXT (IBA LifeSceinces), and 10% glycerol). Eluted proteins were incubated with TEV protease ON at 4 deg C to cleave Strep-GFP tags. Cleaved proteins were subjected to Heparin HP column (5mL) to separate GFP and other non-DNA binding proteins. Peak elutions were pooled and concentrated for subsequent size exclusion chromatography. Concentrated fractions were injected into a Superose 6 increase 10/300 equilibrated with 25mM HEPES pH 7.9 at 4°C, 150 mM NaCl, 2 mM MgCl_2_, 1 mM TCEP, and 10% glycerol). Peak fractions were analyzed by SDS-PAGE and flash frozen in independent fractions and stored at -80 °C until further use.

### Synthesis of peptide compounds

All reactions except those carried out in aqueous phase were performed under an inert atmosphere of argon or nitrogen, unless otherwise noted. Materials were purchased from commercial suppliers and used without further purification, unless otherwise noted. Isolated yields were calculated by weighing products. The weight of the starting materials and the products were not calibrated. Analytical thin layer chromatography (TLC) was performed on Merck silica gel 60 F_254_ plates. Flash column chromatography was performed Biotage Isolera Prime using a SNAP cartridge (Biotage). ^1^H NMR were measured in CD_3_OD solution, and reported in parts per million (ppm) relative to tetramethylsilane (0.00 ppm) as internal standard or referenced to residual solvent peaks of CD_3_OD (3.31 ppm) using JEOL NMM-EC500, JNM-ECX400P or JNM-ECX400, unless otherwise noted. Coupling constant (*J*) was reported in hertz (Hz). Abbreviations of multiplicity were as follows; s: singlet, d: doublet, t: triplet, q: quartet, m: multiplet, br: broad. Data were presented as follows; chemical shift (multiplicity, integration, coupling constant). Mass spectra were obtained on Waters SQ Detector2.

### Cryo-EM sample preparation

Quantifoil Au 2/1 holey carbon grids were cleaned first with chloroform and ethanol. A thin film of carbon (1-2 nm) was deposited on these grids using an in-house carbon floater and the grids were stored for 24 hours at RT before use. Thin carbon coated grids were glow discharged using Tergeo EM plasma cleaner and used immediately. PRC2-activator complex was prepared by mixing ∼4uM PRC2 with 50µM activator in buffer containing 80uM SAH and 0.5mM bis(sulfosuccinimidyl) suberate (BS3). The reaction was quenched with 25mM Tris-base. A tube was prepared that contained 1ul 0.1% Octyl-beta-Glucoside, 8ul of buffer containing 25mM HEPES, 50mM Nacl and no glycerol. 1uL of crosslinked PRC2-cyclopeptide complex was then added and 4ul of the sample was applied to freshly glow dishared carbon coated grids. The grid was incubated for 30 sec in a Vitrobot Mark IV humidified chamber, then blotted for 3-4 s at 5°C and 100% humidity and then plunged into liquid ethane.

### Cryo-EM data collection and processing

Cryo-EM dataset was collected at Pacific Northwest Cryo-EM Center (PNCC) on a Titan Krios equipped with Gatan K3 direct detector and a BioContinuum energy filter. A filter slit width of 20-eV was used for the duration of the collection. Movies were recorded in super resolution mode at a nominal magnification of 130,000x, corresponding to a calibrated pixel size of 0.825Å/pixel (super-resolution 0.4125 Å/pixel). SerialEM was used for automated data acquisition with a defocus range of −2.5 to −0.8 µm. A total dose of 50 electrons per square angstrom (e^−^/Å^2^) fractionated roughly 1 e^-^/A^2^/frame was used for collection. Data was acquired as dark-subtracted, non–gain corrected compressed movies (tif format), and gain correction was applied during motion correction using CryoSparc {Zheng, 2017 #69}.Data was processed in CryoSparc and RELION 4 {Kimanius, 2021 #88}. The movie frames were aligned using CryoSparc and CTF parameters were fit using CryoSparc {Rohou, 2015 #72}. Only micrographs with 4 Å or better CTF fit were retained for further analysis. Initial round of 2D classification within CryoSparc was utilized to remove obvious junk particles. Initial models were generated within CryoSparc using ab initio reconstruction from a subset of the data. Subsequent processing steps including several runs of heterogeneous refinement, homogeneous refinement, particle subtraction and 3D classification, and 3D variability analysis within CryoSparc. The final set of homogeneous particles (665,173) were imported into RELION (using pyem and reliosparc; insert github links) for 3D refinement and the RELION 3D refinement map was imported back into CryoSparc for local resolution filtering. All homogeneous refinements in CryoSparc and RELION used soft-edged masks that were generated within RELION.

### Model Building

Individual PRC2 subunits were built using cryo-EM maps from the final consensus refinement from RELION and local resolution filter map from CryoSparc. The coordinates of nucleosome-bound PRC2 six-subunit complex (PDB: 6WKR) {Kasinath, 2021 #13} was used as a starting model from which all the coordinates were adjusted and rebuilt in the new map using COOT {Emsley, 2004 #74}. All PRC2 subunit models were flexibly fit using ISOLDE to adjust rotamers and outliers. To generate an atomic model of the cyclic peptide activator, the SMILES string corresponding to the chemical structure of the Phe-type activator was used to generate an initial model and restraints from Phenix. This initial model was rigid body docked into the cryo-EM map in ChimeraX followed by flexible fitting and real space refinement in ISOLDE. The resulting coordinates were merged with PRC2 model from ISOLDE to produce a coordinate file containing both PRC2 and ISOLDE. This merged coordinate was subjected to additional rounds of ISOLDE and Phenix real space refinement. The final model demonstrates excellent statistics as analyzed by MolProbity and CaBLAM reports. The map vs model FSC was generated within Phenix and showed excellent consensus. All model and map figures were generated using Chimera and ChimeraX.

### Nucleosome Reconstitution

Xenopus histones were either purified in-house according to (Dyer et al.) or purchased from the Histone Source (Colorado State University). Equimolar ratios of individual H2A, H2B, H3 and H4 were mixed in unfolding buffer and ran on a Superdex 200. Peak fractions were pooled and flash frozen supplemented with 20% glycerol. Mononucleosomes were prepared according to Dyer et al., however a 227bp DNA fragment containing linker sequences was used as nucleosome DNA. Nucleosomes were stored at 4 °C in a buffer containing MES, pH =6.7.

### PRC2-core activity assay

The enzyme mix [55 ng/well PRC2 (active motif #31887) in 18 μL of enzyme buffer (50 mM Tris-HCl pH 8.5, 5 mM DTT, 0.05% Tween20)] was added to each concentration of the sample (DMSO solution, 2 μL) on a 96-well plate and the mixture was incubated for 10 min. The substrate mix {120 nM H3[21-44]-GK biotin (anaspec #AS-64440-025), 3.04 µM SAM and 0.5 μM [^3^H]-SAM (PerkinElmer #NET155V250UC, total 9.25 MBq) in 20 μL enzyme buffer} was added to the solution and the mixture was incubated for 60 min. The enzyme reaction was quenched by adding 50 μL of SAM solution (1 mM). Then, a suspension of Streptavidin PVT SPA Scintillation Beads in a buffer (50 mM Tris-HCl pH 8.5, 0.05% Tween20) (25 mg/mL) was added to the solution. After mixing the suspension, the mixtures on the plate were incubated for 90 min and the CPM counts were measured using MicroBeta (PerkinElmer). This assay was conducted at least twice. When mononucleosomes were used as a substrate, 0.5 µg/well biotinylated mononucleosomes (active motif #31467) ere added to the solution instead of H3[21-44]-GK biotin. All other conditions were performed as described above.

### Western blot activity assays

For all methyltransferase assays using PRC2_119-450_ or PRC2_ApJ 1-450_ was used at a concentration of 200 nM. Mononucleosomes or pure histone H3 were used at 400 nM or 1 µM respectively. Methyltransferase reactions were carried in histone methyltransferase buffer (40uM S-adenosyl methionine, 25 mM HEPES, 50 mM NaCl, 2 mM MgCl_2_ and 1 mM DTT). Either compound #5 or #11 were resuspended in methyltransferase buffer at 10 mM and diluted to a final concentration of 0.1 µM to 200 µM. Reactions took place at room temperature for 90 minutes and quenched with 4X Laemmli buffer. Samples were run on 18% acrylamide gels and transferred to Amersham 0.45 µm membrane (Cytiva) at 4C. Immunoblotting was carried out using antibodies specific for H3K27me1, H3K27me2 and H3K27me3 (Cell Signaling Technology). Detection was performed with a chemiluminescent HRP substrate (Cayman chemicals).

### Culturing and Transfection of mESCs

E14 mESCs were cultured with MEFs supplemented with mLIF. Cells were fed every 24 hours. Tazemetostat was added to the media in DMSO at 10 µM for 7 days and added to the media during passaging. After 7 days, mESCs were washed twice with media to allow Tazemetostat to diffuse out of the cells. After Tazemetostat withdrawal, cyclo-phe (compound #11 in DMSO) was transfected in 1X proteocarry for 1 hour and was subsequently washed away and replenished with new media. Cells in a 6-well dish were collected at specified time points, spun down, and flash-frozen for further analysis.

### Histone extraction from mESCs

mESC cell pellets were lysed in hypotonic buffer (25mM HEPES pH =7.9, 10mM KCl, 0.1% NP-40, 1mM TCEP, 1mM EDTA, 1x PIC) and nuclei were spun down at 9,800xg for 15mins at 4C. The nuclei pellet was then resuspended in 1X RIPA buffer (25mM Tris-HCl, 150mM NaCl, 1mM EDTA, 1% SDS, 0.5% Sodium deoxycholate, 1:1000 benzonase) and sonicated at 25% amplitude in 3 pulses. Nuclear debris was spun down at 15,000xg for 15 mins, and the solubilized nuclear fraction was measured using BCA absorbance, aliquoted, and flash frozen. 2.5 µg total of solubilized pure chromatin was loaded onto an 18% SDS gel and separated. Transfer of histones onto a nitrocellulose membrane for 90V for 10min, and 60V for 30 mins. Primary antibodies (Cell Signaling Tech) were added at 1:5000 overnight. secondary antibodies at 1:7500 for 1hr at room temp. Imaged using chemiluminescence.

### Flow Cytometry

HEK 293 cells adapted to suspension were used. Cyclopeptide transfection was performed using ProteoCarry (Funakoshi) at a concentration of 50 µM. Cells remained in suspension for 5hrs and were washed with fresh medium. Following 48hrs, cells were spun down and resuspended in FACS buffer containing FBS and EDTA. Cells were analyzed and sorted with Aria Fusion (BD Biosciences) and FITC+ cells were collected in FACS buffer. Cells were spun down at 6000rpm and resuspended in 1X SDS-Laemmli buffer.

### Mass Spectrometry for mouse plasma assays

Peptide detection was performed on a Shimadzu Nexera series along with a Shimadzu LCMS6080. Samples were passed over a Acquity UPLC HSS T3 1.8 µm column (3x50 mm) at a flow rate of 0.5 mL/min. Column temperature was maintained at 45 °C. The mobile phase (A) was 0.1 % formic acid/water and mobile phase (B) was 0.1 % formic acid/methanol. Samples were eluted with a gradient: 0 to 0.2 min; 10 % B, 0.2 min – 1 min; 10-90% B, 1-2.55 min; 90% B, 2.55-2.56 min; 90%-10%B; 2.56-4.5 min; 10%B. All assays were performed in two independent experiments.

## Acknowledgments

We thank Erik Hartwick, Garry Morgan for EM support, Theresa Nahreini for assistance with flow cytometry, and Annette Erbse for assistance with shared instrument facility at CU Boulder Biochemistry. We thank Vamseedhar Rayaprolu and the Pacific Northwest Cryo-EM center (PNCC) for EM data collection. We thank Ms. Hiroko Azuma (Hokkaido University) for assistance of immunoblotting analysis using core PRC2.

## Funding

This work was founded in part by JSPS KAKENHI Grant-in-Aid for Scientific Research (C) (19K06988), the Japan Agency for Medical Research and Development (AMED) under Grant Number 18ae0101047h0001, 19ae0101047h0002 and 20ae0101047h0003, Astellas Foundation for Research on Metabolic Disorders, The Uehara Memorial Foundation, Kobayashi foundation for cancer research and Suhara memorial foundation to F.Y. This work was partially funded by NIH grants GM132544 and GM155426 to V.K. This work was founded in part by Grant-in-Aid for JSPS Fellows (20J21033) to Y.T. This work was supported by the Hokkaido University Global Facility Center, the Pharma Science Open Unit (PSOU), funded by MEXT under “Support Program for Implementation of New Equipment Sharing System”, and the Platform Project for Supporting Drug Discovery and Life Science Research (Basis for Supporting Innovative Drug Discovery and Life Science Research; BINDS) from AMED under Grant Number JP24ama121053. We thank the shared instrument pool (RRID: SCR_018986), Department of Biochemistry, University of Colorado, Boulder for the use of the Typhoon 5 imager and Beckman Avanti centrifuges. Typhoon 5 was funded by NIH Grant S10OD034218-01 and the Beckman centrifuges by NIH Grant R24OD033699-01.

## Author contributions

Y.T and F.Y. designed compounds. Y.T., A.I., S.Y. and T.H. synthesized compounds. Y.T. performed conformational analysis of the synthesized compounds. F.Y. organized and established experimental procedures and performed *in vitro* biochemical activity assays using core PRC2. M.M established experimental procedures, expressed and purified different PRC2 complexes and nucleosomes, prepared the samples and cryo-grids, collected and processed the EM data, carried out model building and refinement, performed all in vitro biochemical activity assays with 6-subunit PRC2, and carried out flow cytometry experiments. A.Z. and R.S. assisted in preparing histones, DNA, and reconstitution of nucleosomes. L.Y. carried out the peptide transfections and flow cytometry experiments with M.M. L.B.P and N.L assisted M.M in constructing the PRC2 mutations, expression and purification of PRC2 containing mutations, and biochemical activity assays. E. O carried out the mESC experiments with advice from M.M and J.B. K.K. and N.N. performed experiments related to the compound stability in mouse plasma. V.K. and F.Y. conceived the project. V.K. analyzed all the cryo-EM data, biochemical assays with 6-subunit PRC2, and flow cytometry data. F.Y. analyzed all the peptide synthesis, biochemical assays with PRC2-core, and mouse plasma stability data. S.I. organized the project. V.K. and F.Y. supervised the entire study. M.M., F.Y., and V.K. wrote the manuscript with input from all authors.

## Competing interests

Authors declare no competing interests.

## Data and materials availability

Cryo-EM density maps, half maps, local resolution filtered maps, masks and fitted models have been deposited in the Electron Microscopy Data Band (EMDB) and the Protein Data Bank (PDB) for the PRC2-AEBP2-JARID2 (aa 119-450) bound to Phe-type activator [EMDB: EMD-70352 and PDB: 9ODA.

## Notes

### Competing Interest Statement

The authors have declared no competing interest.

### Summary of Updates

The manuscript files were not changed. In the original submission, corresponding author was not designated. This was revised.

