## Supplemental Information for "Structure-guided design and development of cyclic peptide allosteric activators of Polycomb Repressive Complex 2"

#### Synthesis of amino acids

**Scheme S1**. Synthesis of Fmoc-Lys(me3)-OH (**S2**)[1)]

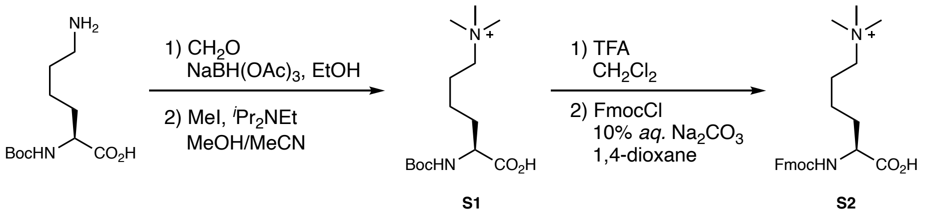


**Boc-Lys(me3)-OH (S1)**

A solution of Boc-Lys-OH (985 mg, 4.00 mmol) in EtOH was treated with 37% *aq.* $CH_2O$ (972 µL, 12.0 mmol) at 0 °C for 10 min. Then, $NaBH(OAc)_3$ (3.38 g, 16.0 mmol) was added to the solution, and the mixture was stirred at room temperature (rt) for 12 h. When the reaction was not completed after 12 h, $NaBH(OAc)_3$ was added as needed. The progress of the reaction was monitored by silica gel TLC with 5% $H_2O$ in MeOH as a mobile phase. After completion, the reaction was quenched with *sat. aq.* $NaHCO_3$. pH of the solution was adjusted to 7, and volatiles were removed *in vacuo*. The residue was dissolved in $H_2O$ (15 mL), and the aqueous phase was extracted with $CHCl_3$ (20 mL×25). The combined organic phase was dried ($Na_2SO_4$), filtered, and concentrated. The residue containing the desired compound was used for the next step without further purification.

A solution of the residue (250 mg, 0.91 mmol) and DIPEA (201 µL, 1.18 mmol) in MeCN/MeOH (2:1, 9 mL) was treated with iodomethane (567 µL, 9.11 mmol) at rt for 4 h. The progress of the reaction was monitored by silica gel TLC with $CHCl_3$/MeOH/AcOH (4:5:1) as a mobile phase. The mixture was concentrated *in vacuo*, and then the residue was dissolved in MeOH/0.1 M *aq.* NaOH (1:1, 18 mL). After stirring at rt for 1 h, the reaction was neutralized with 1 M *aq.* HCl, and the mixture was concentrated *in vacuo*. The residue was purified by silica gel column chromatography (ϕ 2 × 4 cm, MeOH/$Et_3N$ = 97:3) to afford **S1** (200 mg, 0.616 mmol, 65% over 2 steps) as a white foam. $^1H$ NMR ($CD_3OD$, 400 MHz) δ 3.98 (m, 1H, α-C*H*), 3.11 (s, 9H, *N*-Me), 1.88-1.64 (m, 4H, β-C*H*, δ-C*H*), 1.43 (s, 11H, γ-C*H*, $^t$Bu), the solvent peaks overlapped with the sample peaks (ε-C*H*); ESIMS-LR *m/z* calcd for $C_{14}H_{29}N_2O_4$ [$M^+$] 289.2, found 289.1.

**Fmoc-Lys(me3)-OH (S2)**

A solution of **S1** (200 mg, 0.616 mmol) in TFA/dichloromethane (1:1, 12 mL) was stirred at rt for 2 h. The progress of the reaction was monitored by silica gel TLC with 5% $H_2O$ in MeOH as a mobile phase. After completion of the reaction, the solvent was removed *in vacuo*. The residue in 1,4-dioxane/10% *aq.* $Na_2CO_3$ (1:1, 12 mL) was treated with FmocCl (175 mg, 0.678 mmol) at rt for 12 h. The progress of the reaction was monitored by silica TLC with 5% $H_2O$ in MeOH as a mobile phase. After completion of the reaction, the mixture was acidified with 1 M *aq.* HCl and the solvent was removed *in vacuo*. The residue was purified by silica gel column chromatography (ϕ 2 × 4 cm, $CHCl_3$/MeOH = 1:1→0:1) to afford **S2** (155 mg, 0.346 mmol, 56% over 2 steps) as a white foam. $^1H$ NMR ($CD_3OD$, 400 MHz) δ 7.80 (d, 2H, Ph-*H*, *J* = 7.8 Hz), 7.65 (d, 2H, Ph-*H*, *J* = 8.7 Hz), 7.41-7.29 (m, 4H, Ph-*H*), 4.40-4.28 (m, 2H), 4.22 (m, 1H), 4.05(m, 1H, α-C*H*), 3.09 (s, 9H, *N*-Me), 1.89-1.70 (m, 4H, β-C*H*, δ-C*H*), 1.46-1.40 (m, 2H, γ-C*H*), the solvent peaks overlapped with the sample peaks (ε-C*H*); ESIMS-LR *m/z* calcd for $C_{24}H_{31}N_2O_4$ [$M^+$] 411.2, found 411.3.

#### Solid phase peptide synthesis

**general procedure**

**G-1: Resin loading**

Each 2-chlorotrityl resin was placed in a 5 mL polypropylene syringe fitted with a polyethylene filter disc. Each resin was agitated with $CH_2Cl_2$ for 1 h. After removal of $CH_2Cl_2$, a solution of Fmoc protected amino acids (3.0 eq.) and DIPEA (8.0 eq.) in $CH_2Cl_2$ was added to the mixture at rt. After agitation for 1 h, the solvent and soluble reagents were removed by suction. The resins were washed with DIPEA/MeOH/$CH_2Cl_2$ (1:2:17, 2 mL×3), DMF (2 mL×3) and $CH_2Cl_2$ (2 mL×3) to afford resins supported Fmoc protected amino acids.

The amount of loading on resins was determined as follows. Dried resins supported Fmoc protected amino acids (approx.

5 mg; The amount used should be accurately recorded.) were stirred with DMF (2 mL) for 30 min, then DBU (40 μL) was added to the mixture. The mixture was stirred for 30 min. The supernatant was diluted with MeCN (7.96 mL). This solution (1 mL) was further diluted with MeCN (11.5 mL) and subjected to UV measurement at 294 nm. The loading rate was determined from the following equation.

$$\textit{Loading (mmol/g)} = A_{294nm}*12.5*10*1000 / (W*8794)$$

W: weight of the resin used (mg)

**G-2-1: Kaiser test (for primary amine detection)**

A small amount of resin (approx. 1 mg) was treated with a drop of reagent A, B, and C (see below). The mixture was refluxed immediately. When the color of the solution turns blue or purple, it indicates the presence of primary amines. No change of the color indicates that no exposed primary amines are present, suggests the completion of the coupling reaction.

Reagent A; 0.50 g of ninhydrin in 10 mL EtOH
Reagent B; 8.0 g of phenol in 2.0 mL EtOH
Reagent C; 0.13 mg of KCN in 10 mL pyridine

**G-2-2: Chloranil test (for secondary amine detection)**

A small amount of resin (approx. 1 mg) was treated with a drop of reagent D and E (see below). The mixture was allowed to stand for 5 min. When the color of the solution turns blue, it indicates the presence of primary or secondary amines. No change of the color indicates that no exposed amines are present, suggests the completion of the coupling reaction.

Reagent D; 2% *p*-chloranil in DMF
Reagent E; 2% acetaldehyde in DMF

**G-3: Fmoc group removal**

The agitated resins were treated with 20% piperidine/DMF for 5 min, then piperidine/DMF for 15 min to remove the Fmoc protecting groups. The resins were washed with DMF (2 mL×3) and $CH_2Cl_2$ (2 mL×3), and Kaiser test or Chloranil test indicated the completion of all deprotection reactions.

**G-4: Coupling reaction**

A solution of Fmoc protected amino acids (4.0 eq.), HBTU or HATU (3.9 eq.) and DIPEA (8.0 eq) in DMF (1 mL) was added to the mixture of resins, which were agitated for 2 h. All the resins were washed with DMF (2 mL×3) and $CH_2Cl_2$ (2 mL×3) to afford resins supported protected peptides. Kaiser test or chloranil test indicated completion of all coupling reactions. The loading rate was determined as described above.

**G-5: Acetylation**

When the reaction was not completed, the remaining free amino groups were capped by treatment with acetic anhydride. The agitated resins were treated with 20% $Ac_2O/CH_2Cl_2$ for 10 min. The resins were washed with $CH_2Cl_2$ (2 mL×4) and Kaiser test indicated completion of the reactions.

**G-6: Cleavage and deprotection (linear peptides)**

Before cleavage of desired liner peptides, the final Fmoc protecting groups were removed following 'the G-2: Fmoc group removal' procedure. Then, the resins were treated with TFA/$H_2O$/$^i Pr_3SiH$ (95:2.5:2.5). The reaction mixture was stirred at rt for 2 h before filtering the resins. The filtrate was concentrated, washed with $Et_2O$, dried *in vacuo*. The residue was purified by RP-HPLC to afford the linear peptides.

**G-7: Cleavage, macrocyclization and deprotection (cyclic peptides)**

Before cleavage of desired liner peptides, the final Fmoc protecting groups were removed following 'the G-2: Fmoc group removal' procedure. Then, the resins were treated with TFA/$CH_2Cl_2$ (1:99). The reaction mixture was stirred at room temperature for 30 min before filtering resins. The filtrate was concentrated, washed with $Et_2O$ and dried *in vacuo* to afford protected liner peptides. A solution of the peptides in DMF (5 mM) was treated with DPPA (3.0 eq.) and $NaHCO_3$ (5.0 eq.) at rt for 24 h. The solution was filtered and DMF was removed *in vacuo*, then the residue was washed with $Et_2O$ and $H_2O$. The residue was dried *in vacuo* to afford a crude of protected cyclic peptides. The protecting groups of the peptides were removed under conditions with TFA/$H_2O$/$^i Pr_3SiH$ (95:2.5:2.5). The reaction mixture was stirred at rt for 2 h. The mixture was concentrated, washed with $Et_2O$ and dried *in vacuo.* The residue was purified by RP-HPLC to afford the cyclic peptides.

**KAARK(me3)SAPA (2)**

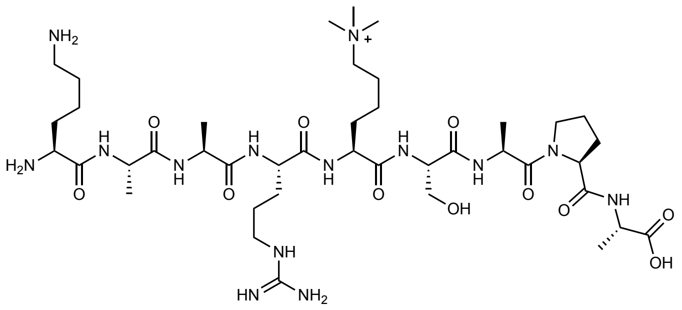

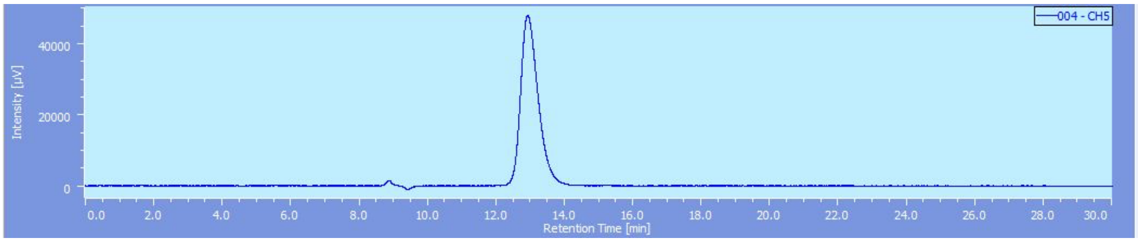


**KAARK(me3)SAP (3)**

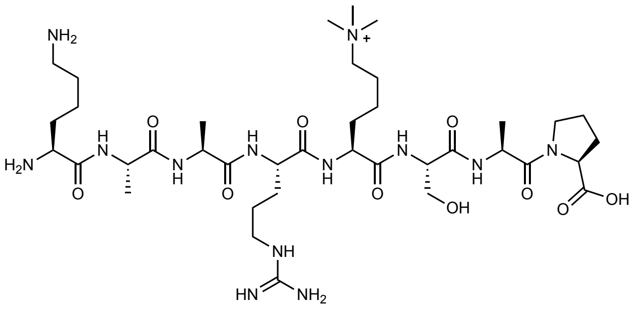

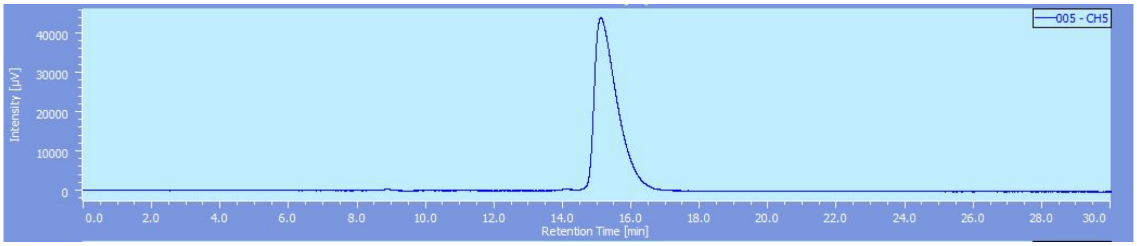


**TKAARK(me3)SA (4)**

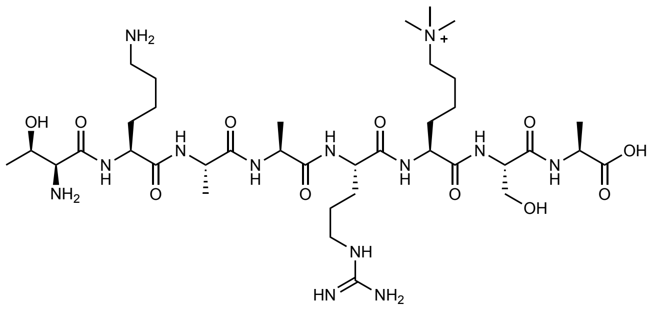

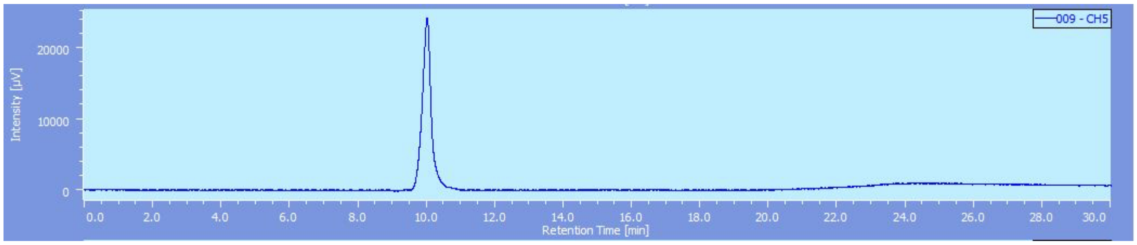


**Supplementary Figure 1. Synthesis of histone H3K27me3 linear peptides.** Compound **1** ([KAARK(me3)SAPATGG**])** was purchased from Anaspec (#AS-64378-1). The overall yield of compounds **2**, **3**, and **4** after HPLC purification (cosmosil $5C_{18}$-PAQ, 250 x 20 mml.D.) was 51%, 63%, and 29%, respectively and the corresponding retention times on the column were 12.9 min, 15.1 min, and 10 min, respectively. Electrospray ionization mass spectrometry confirmed the identities of each compound. The predicted and observed ESIMS-LR m/z matched well for each compound (compound **2** - ($C_{41}H_{78}N_{14}O_{11}$ $[M+H]^{2+}$) predicted 471.3 and observed 471.3; compound **3** - $C_{38}H_{73}N_{13}O_{10}$ $[M+H]^{2+}$ predicted 435.8 and observed 435.7; compound **4** - $C_{37}H_{73}N_{13}O_{11}$ $[M+H]^{2+}$ predicted 437.8 and observed 437.7). The predicted and observed ESIMS-HR m/z also matched well for each compound (compound **2** - ($C_{41}H_{77}N_{14}O_{11}$ $[M]^{+}$) predicted 941.5891 and observed 941.5866; compound **3** - $C_{38}H_{72}N_{13}O_{10}$ $[M]^{+}$ predicted 870.5520 and observed 870.5501; compound **4** - $C_{37}H_{73}N_{13}O_{11}$ $[M+H]^{2+}$ predicted 437.7771 and observed 437.7765).

***cyclo*[KAARK(me3)SAP] (5)**

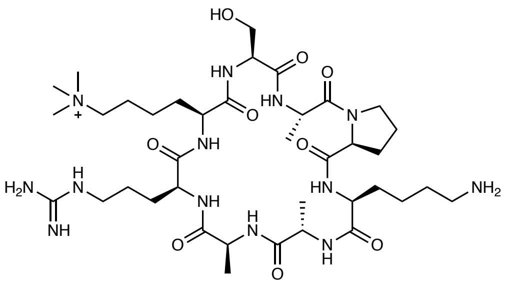

overall yield 43%, $^1$H NMR ($CD_3OD$, 400 MHz) δ 8.92 (br, 1H, amide-N*H*), 8.52 (br, 1H, amide-N*H*), 8.26 (d, 1H, Ala-N*H*, *J* = 9.6 Hz), 8.08 (d, 1H, Ala-N*H J* = 6.4 Hz), 7.37 (m, 2H, Arg-guanidino-N*H*), 4.69 (m, 1H, Ala-α-C*H*), 4.55 (m, 1H, Ala-α-C*H*), 4.41 (t, 1H, Pro-α-C*H*, *J* = 8.2 Hz), 4.20 (m, 1H, α-C*H*), 4.15 (m, 1H, Ser-α-C*H*), 4.03 (m, 1H, α-C*H*), 3.80-3.95 (m, 3H, Ala-α-C*H*, Ser-β-C*H*), 3.71-3.80 (m, 1H, Pro-δ-C*H*), 3.57-3.68 (m, 1H, Pro-δ-C*H*), 3.12 (s, 9H, $NMe_3$), 2.93 (t, 2H, Lys-ε-C*H*, J = 7.8 Hz), 2.29-2.40 (m, 1H, Pro-β-C*H*), 1.19-2.11 (m, 19H), 1.45 (d, 3H, Ala-β-C*H*, *J* = 6.4 Hz), 1.37 (d, 3H, Ala-β-C*H*, *J* = 6.4 Hz), 1.23 (d, 3H, Ala-β-C*H*, *J* = 6.4 Hz), the solvent peaks overlapped with the sample peaks (α-C*H*, Arg-δ-C*H*, Lys(me3)-ε-C*H*, determined by $^1$H-$^1$H COSY); ESIMS-LR *m/z* found 426.5 $[M+H]^{2+}$; ESIMS-HR *m/z* calcd. for $C_{38}H_{70}N_{13}O_9$ $[M]^+$ 852.5414, found 852.5382.

HPLC chart at 220 nm, after purification (cosmosil $5C_{18}$-PAQ, 250 x 20 mmI.D., A: 11.5%, B: 88.5%, $t_r$ = 7.4 min).

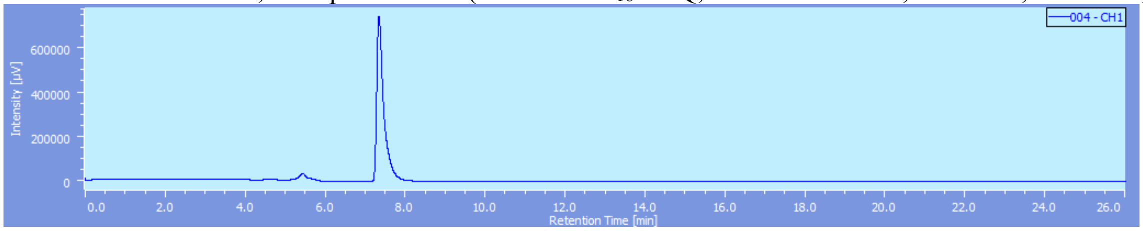


***cyclo*[KAARASAP] (6)**

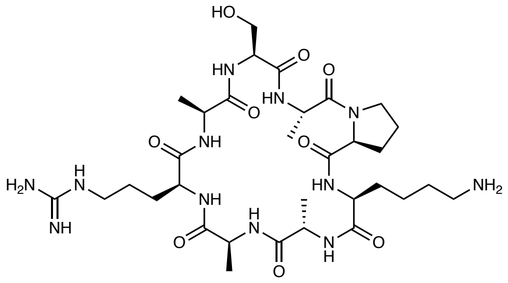

overall yield 23%, ESIMS-LR *m/z* calcd.for $C_{32}H_{56}N_{12}O_9$ $[M]^+$ 752.4, found 752.4; ESIMS-HR *m/z* calcd. for $C_{32}H_{57}N_{12}O_9$ $[M+H]^+$ 753.4366, found 753.4379.

HPLC chart at 220 nm, after purification (cosmosil $5C_{18}$-PAQ, 250 x 20 mmI.D., A: 10%, B: 90%, $t_r$ = 24.3 min).

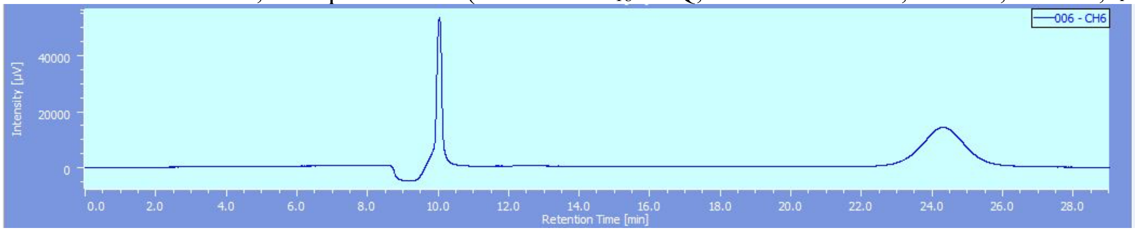


***cyclo*[KAARK(me3)SAA] (7)**

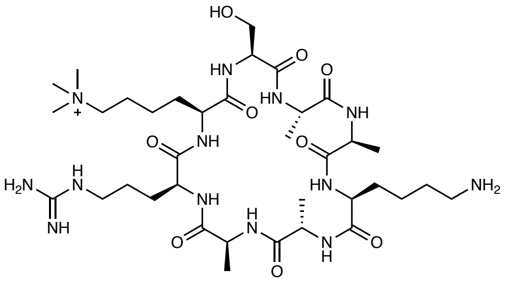

overall yield 7%; ESIMS-HR *m/z* calcd. for $C_{36}H_{68}N_{13}O_9$ $[M]^+$ 826.5257, found 826.5256.

HPLC chart at 220 nm, after purification (cosmosil $5C_{18}$-PAQ, 250 x 20 mmI.D., A: x%, B: x%, $t_r$ = x min).

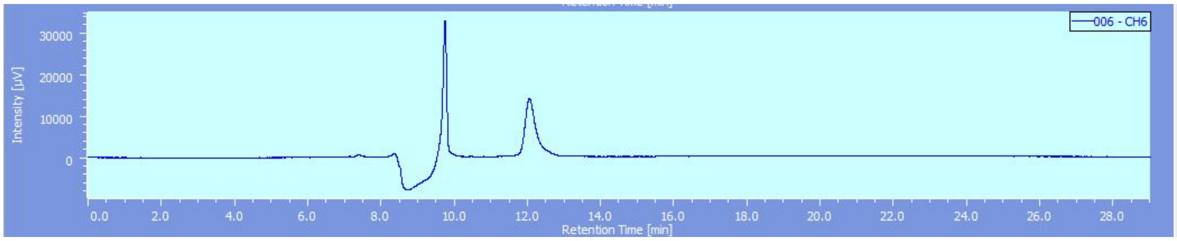


##### *cyclo*[AAARK(me3)SAP] (8)

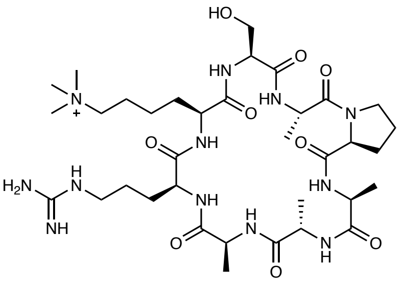


overall yield 18%, ESIMS-LR *m/z* calcd. for $C_{35}H_{63}N_{12}O_9$ $[M+H]^{2+}$ 398.2, found 398.2; ESIMS-HR *m/z* calcd. for $C_{35}H_{63}N_{12}O_9$ $[M]^+$ 795.4835, found 795.4857.

HPLC chart at 220 nm, after purification (cosmosil $5C_{18}$-PAQ, 250 x 20 mmI.D., A: 13%, B: 87%, $t_r$ = 15.5 min).

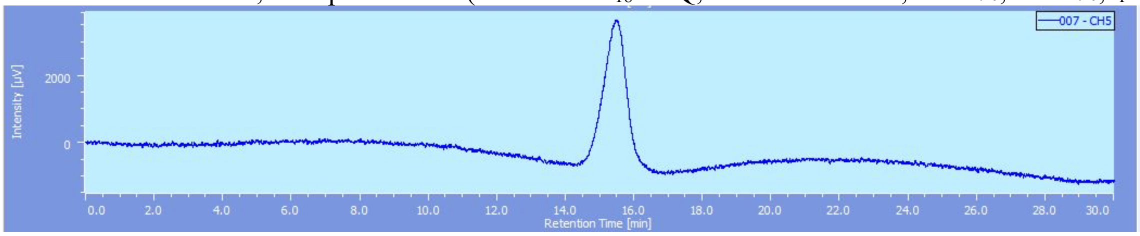


##### *cyclo*[KAAAK(me3)SAP] (9)

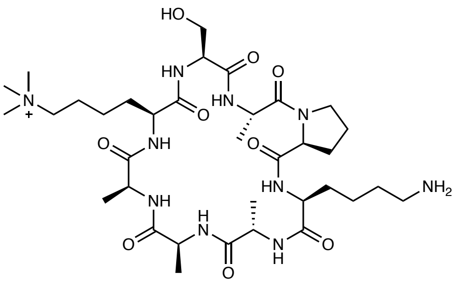


overall yield 26%, ESIMS-LR *m/z* calcd. for $C_{35}H_{63}N_{10}O_9$ $[M]^{2+}$ 767.5, found 767.5; ESIMS-HR *m/z* calcd. for $C_{35}H_{63}N_{10}O_9$ $[M]^+$ 767.4774, found 767.4757.

HPLC chart at 220 nm, after purification (cosmosil $5C_{18}$-PAQ, 250 x 20 mmI.D., A: 10%, B: 90%, $t_r$ = 13.8 min).

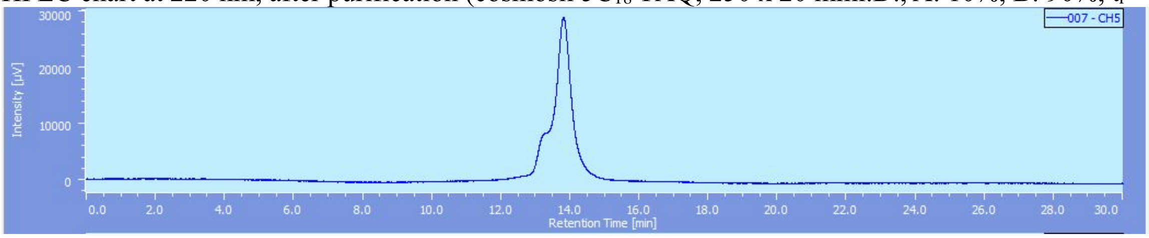


##### *cyclo*[AAAAK(me3)SAP] (10)

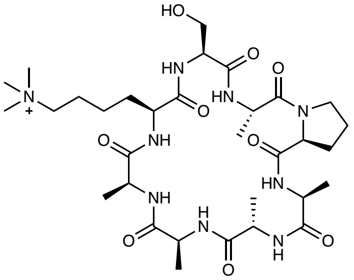


overall yield 10%, ESIMS-LR *m/z* calcd. for $C_{32}H_{56}N_9O_9$ $[M]^+$ 710.4, found 710.5; ESIMS-HR *m/z* calcd. for $C_{32}H_{56}N_9O_9$ $[M]^+$ 710.4196, found 710.4163.

HPLC chart at 220 nm, after purification (cosmosil 5C$_{18}$-PAQ, 250 x 20 mmI.D., A: 15%, B: 85%, $t_r$ = 11.4 min)

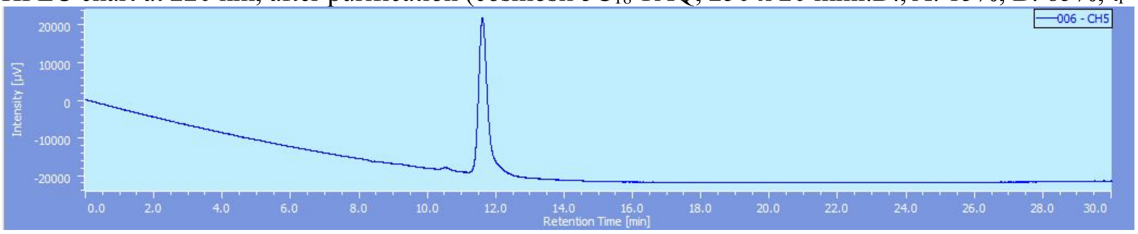


***cyclo*[KAARK(me3)FAP] (11)**

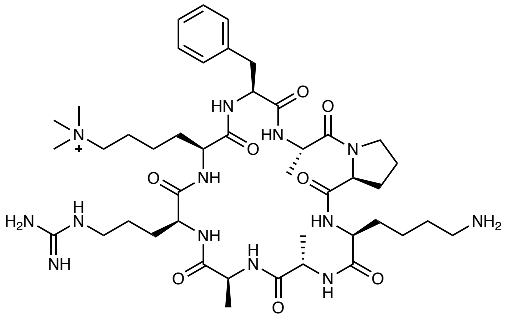

overall yield 30%, $^1$H NMR (CD$_3$OD, 400 MHz) δ 8.79 (br, 1H, amide-N*H*), 8.60 (d, 1H, amide-N*H*, *J* = 4.6 Hz), 8.02-8.08 (m, 2H, Ala-amide-N*H*), 7.58 (br, 1H, amide-N*H*), 7.50 (t, 1H, *J* = 6.0 Hz), 7.26-7.35 (m, 6H, Ph-*H*, Arg-guanidino-N*H*), 4.76 (m, 1H, Ala-α-C*H*), 4.69 (t, 1H, α-C*H*, *J* = 8.7 Hz), 4.48-4.58 (m, 2H, Ala-α-C*H*, Phe-α-C*H*), 4.38 (t, 1H, Pro-α-C*H*, *J* = 7.8 Hz), 4.13-4.20 (m, 1H, α-C*H*), 3.88 (t, 1H, Ala-α-C*H*, *J* = 6.8 Hz), 3.72-3.82 (m, 2H, α-C*H*, Pro-δ-C*H*), 3.53-3.60 (m, 1H, Pro-δ-C*H*), 3.08-3.15 (m, 11H, NMe$_3$, Arg-δ-C*H*), 2.95 (t, 2H, Lys-ε-C*H*, *J* = 7.3 Hz), 2.29-2.38 (m, 1H, Pro-β-C*H*), 1.12-2.11 (m, 19H), 1.46 (d, 3H, Ala-β-C*H*, *J* = 6.4 Hz), 1.38 (d, 3H, Ala-β-C*H*, *J* = 6.4 Hz), 1.24 (d, 3H, Ala-β-C*H*, *J* = 6.4 Hz), the solvent peaks overlapped with the sample peaks (Lys(me3)-ε-C*H*, Phe-β-C*H*, determined by $^1$H-$^1$H COSY); ESIMS-LR *m/z* found 456.5 $[M+H]^{2+}$; ESIMS-HR *m/z* calcd. for $C_{44}H_{74}N_{13}O_8$ $[M^+]$ 912.5778, found 912.5777.

HPLC chart at 220 nm, after purification (cosmosil 5C$_{18}$-PAQ, 250 x 20 mmI.D., A: 15%, B: 85%, $t_r$ = 24.7 min).

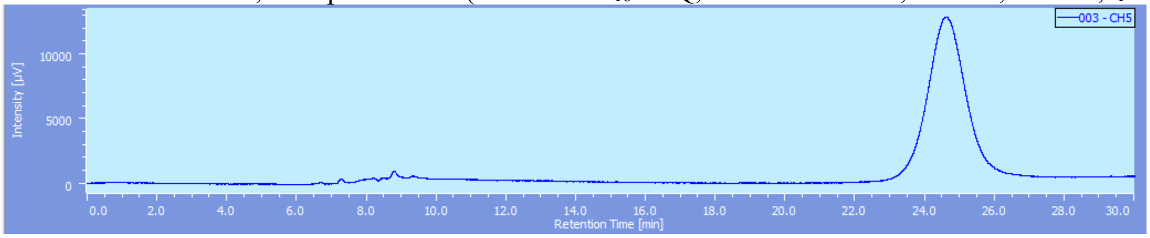


***cyclo*[KAARK(me3)YAP] (12)**

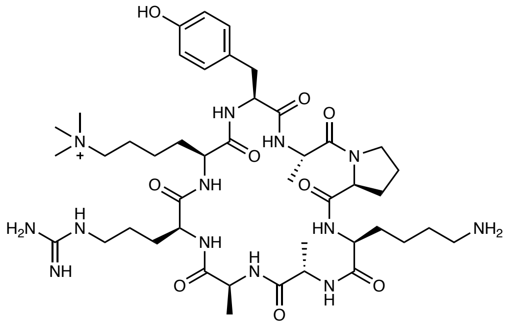

overall yield 23%, ESIMS-LR *m/z* calcd. for $C_{44}H_{74}N_{13}O_9$ $[M+H]^{2+}$ 464.8, found 464.8; ESIMS-HR *m/z* calcd. for $C_{44}H_{74}N_{13}O_9$ $[M]^+$ 928.5727, found 928.5712.

HPLC chart at 220 nm, after purification (cosmosil 5C$_{18}$-PAQ, 250 x 20 mmI.D., A: 15%, B: 85%, $t_r$ = 29.6 min).

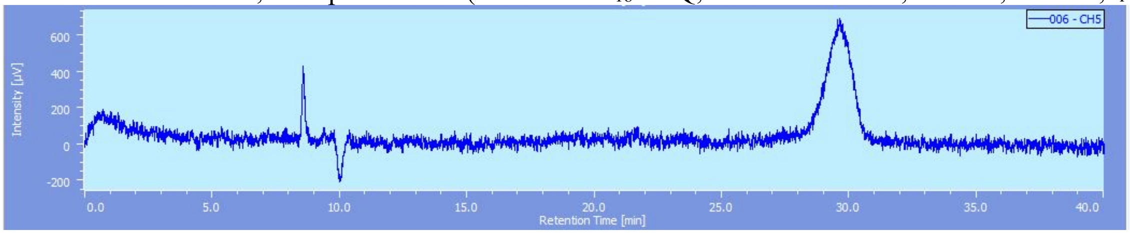


***cyclo*[KAARK(me3)WAP] (13)**

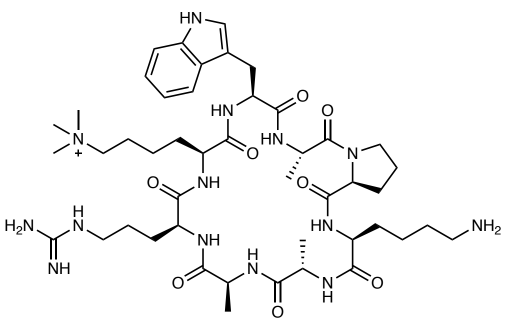

overall yield 38%, ESIMS-LR *m/z* calcd. for $C_{46}H_{76}N_{14}O_8$ $[M+H]^{2+}$ 476.3, found 476.3; ESIMS-HR *m/z* calcd. for $C_{46}H_{75}N_{14}O_8$ $[M]^+$ 751.5887, found 751.5870.

HPLC chart at 220 nm, after purification (cosmosil $5C_{18}$-PAQ, 250 x 20 mmI.D., A: 19%, B: 81%, $t_r$ = 13.7 min).

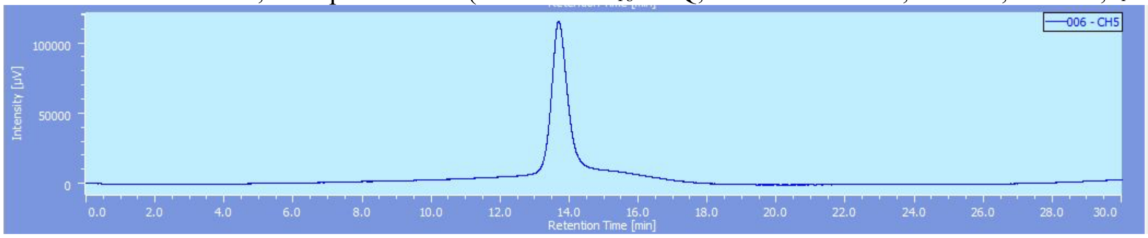


***cyclo*[K(FITC)AARK(me3)FAP] (14)**

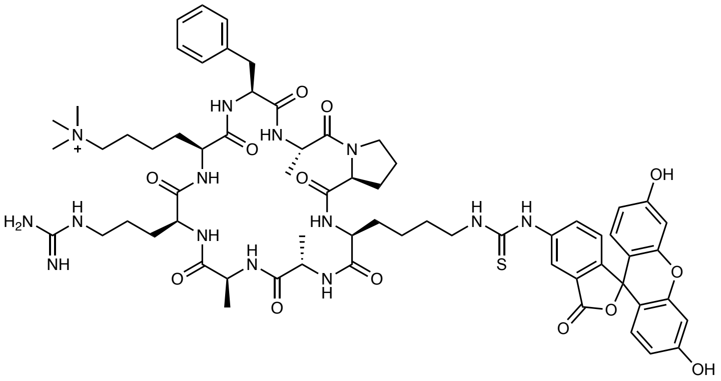

A solution of **11** (10 mg, 11.0 µmol) and $^i$Pr$_2$NEt in DMF (500 µL) was treated with FITC (4.3 mg, 11.0 µmol) at room temperature for 2 h in the dark. The progress of the reaction was monitored by ESI-MS. After completion of the reaction, solvents were removed in vacuo. The residue was purified by RP-HPLC to afford 14 (9.1 mg, 64%) as yellow solid.
ESIMS-HR *m/z* calcd. for $C_{65}H_{85}N_{14}O_{13}S$ $[M]^+$ 1301.6136, found 1301.6088.

HPLC chart at 220 nm, after purification (cosmosil $5C_{18}$-PAQ, 250 x 20 mmI.D., A: 33%, B: 67%, $t_r$ = 17.5 min).

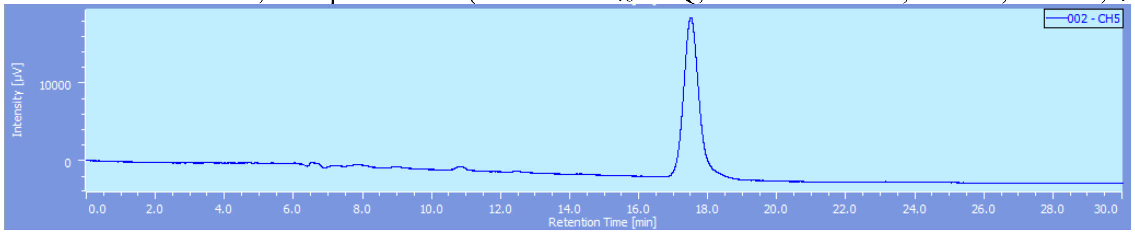


**Supplementary Figure 2**. Synthesis, yield, and mass spectrometry validation of all cyclic peptides (compounds **5-14**)

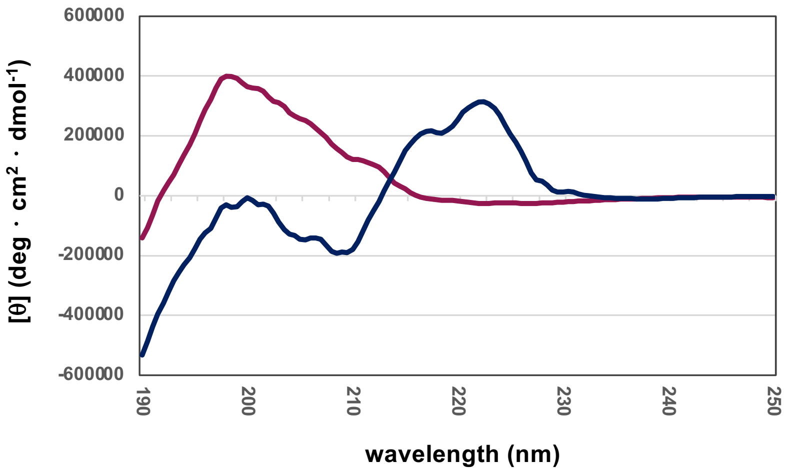


**Supplementary Figure 3**. Circular dichroism spectra of Ser-type cyclic peptide (compound **5**; *cyclo*[KAARK(me3)SAP]; shown in red) and linear peptide (compound **3**; KAARK(me3)SAP; shown in blue).

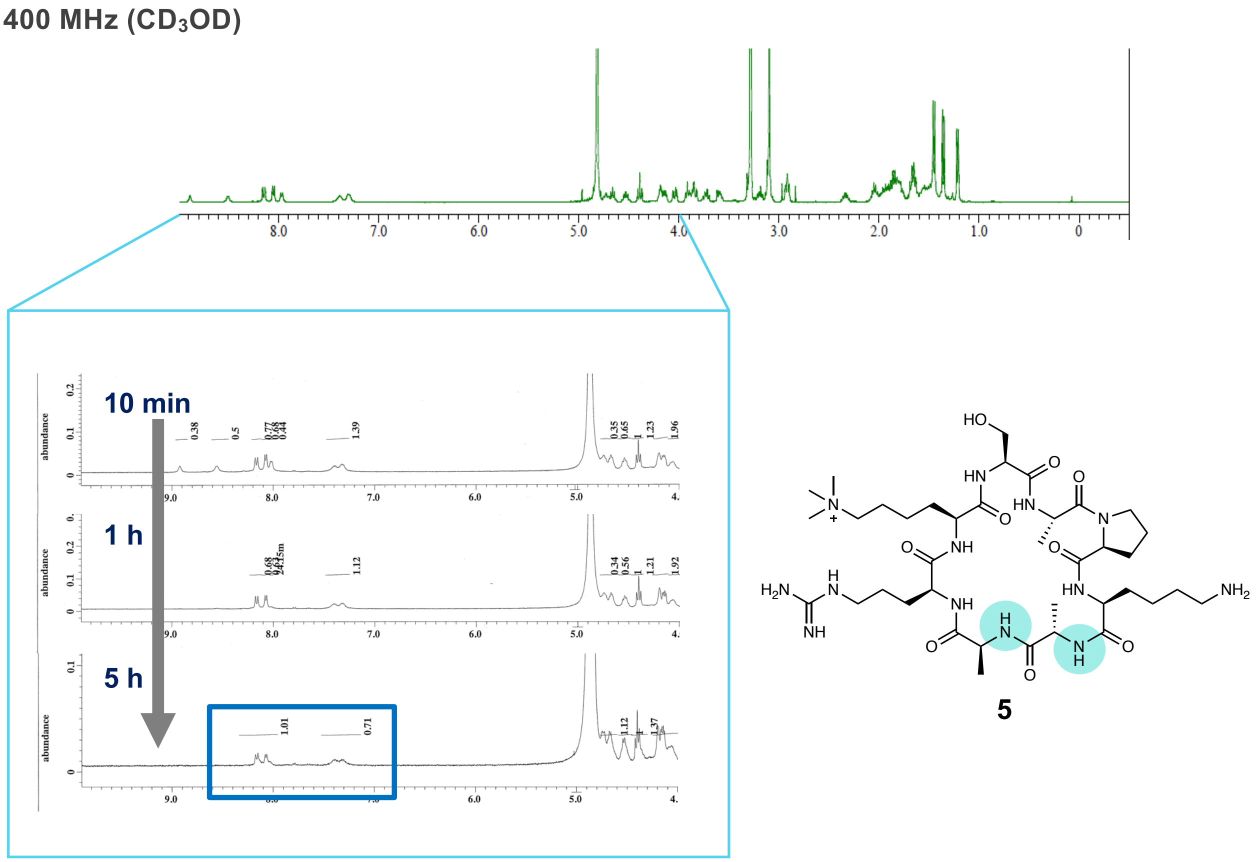


**Supplementary Figure 4**. [1]H NMR chart of Ser-type cyclic peptide (compound **5**). Two Ala residues of compound **5** form intramolecular hydrogen bonds.

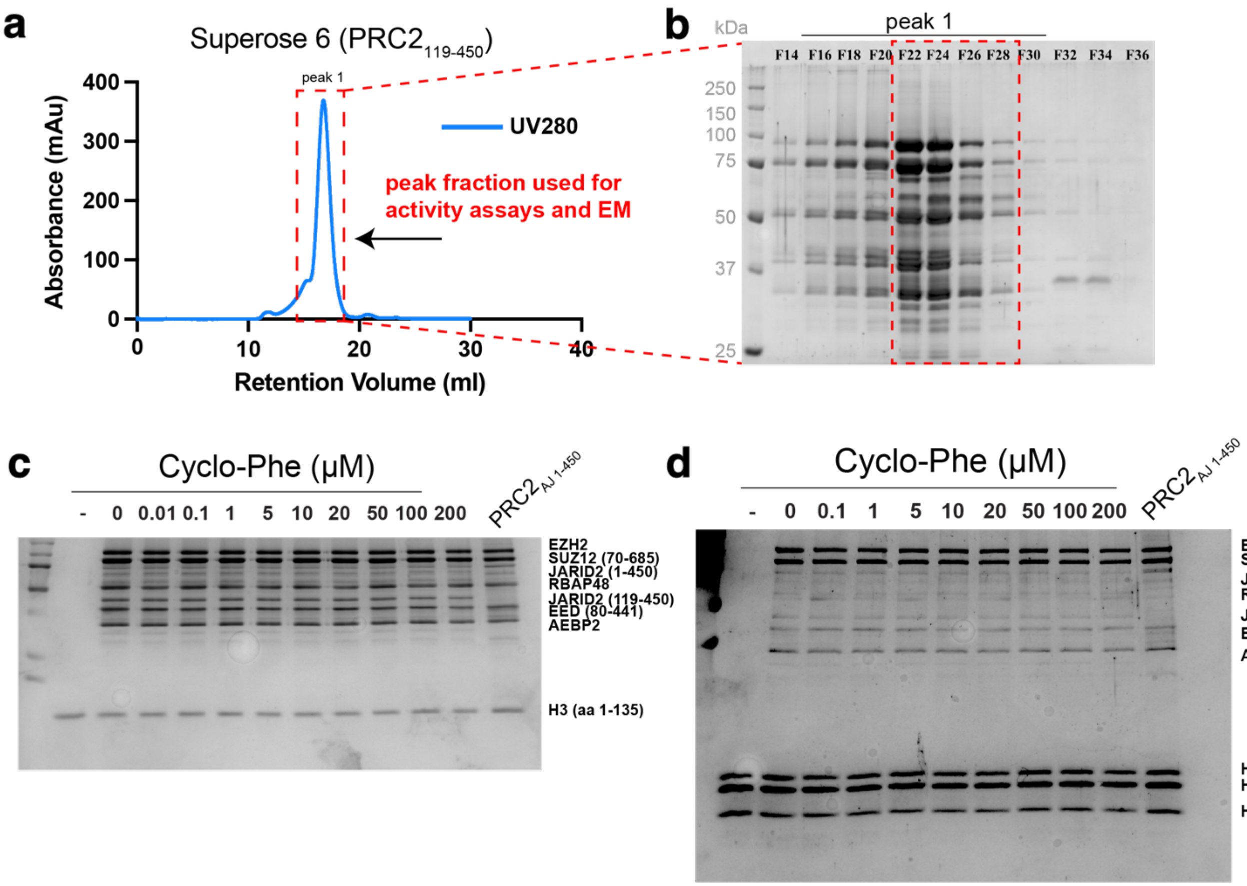


**Supplemental Figure 5. Purification of PRC2-AJ$_{119–450}$ and corresponding Coomassie stained gels for methyltransferase activity assays. (A-B)** Size exclusion chromatogram and corresponding SDS-PAGE gel of recombinantly expressed PRC2 $_{AJ119-560}$ fractions used in either EM studies or activity assays. **(C)** Representative Coomassie stained SDS-PAGE gel for activity assay on recombinant histone H3. **(D)** Representative Coomassie stained SDS-PAGE gel for activity assay on recombinant mononucleosomes.

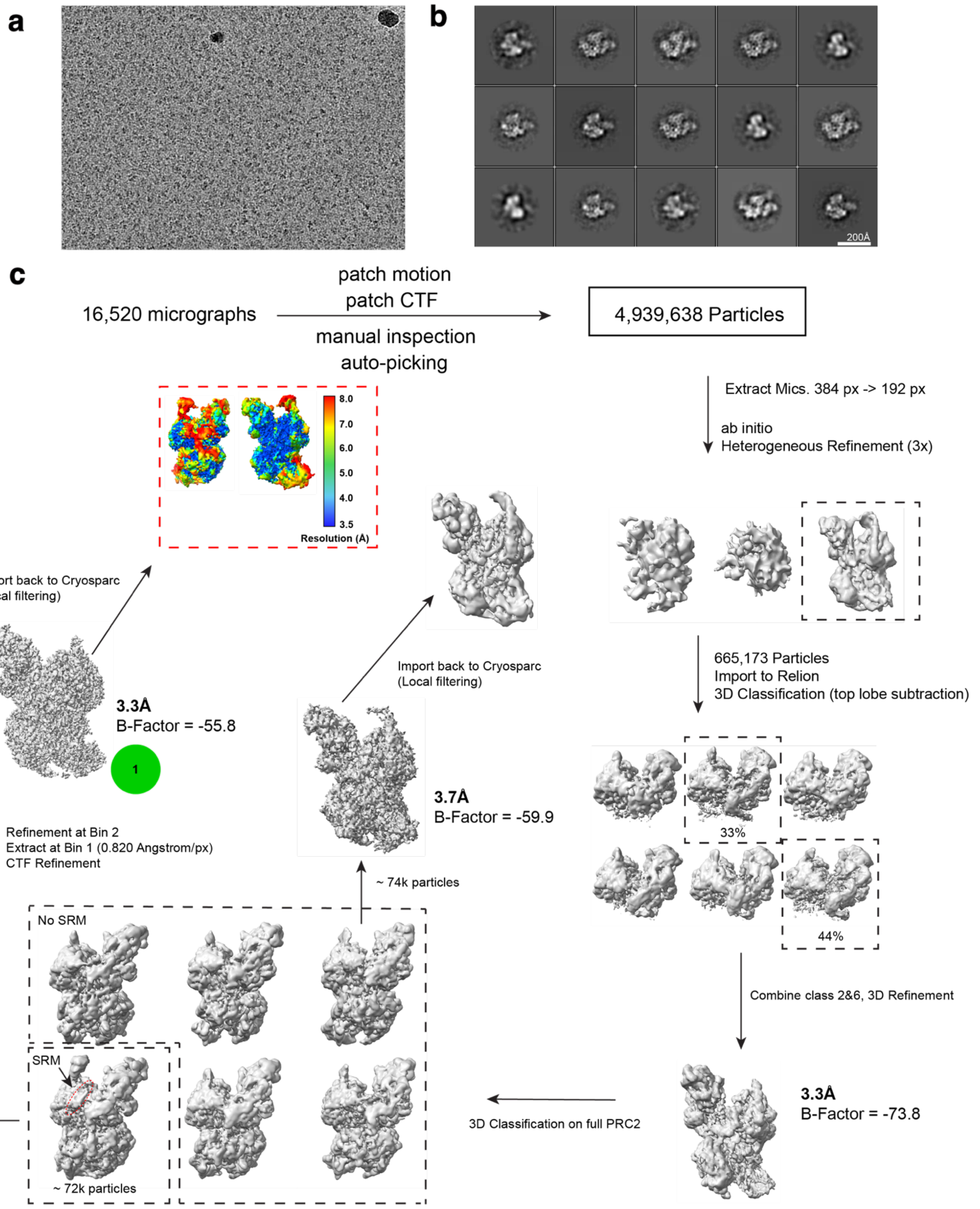


**Supplemental Figure 6. Structure determination of the PRC2 $AJ_{119\text{-}450}$ cyclopeptide complex. (A)** Representative micrograph for sample and grid collected. **(B)** Representative RELION-generated 2D classes for subset of particles used in final reconstruction. **(C)** Workflow of cryo-EM data processing. Final map used for reconstruction was imported back into Cryosparc for local filtering. Range of resolution from 3.5 Å - 8 Å is depicted for final map used for model building.

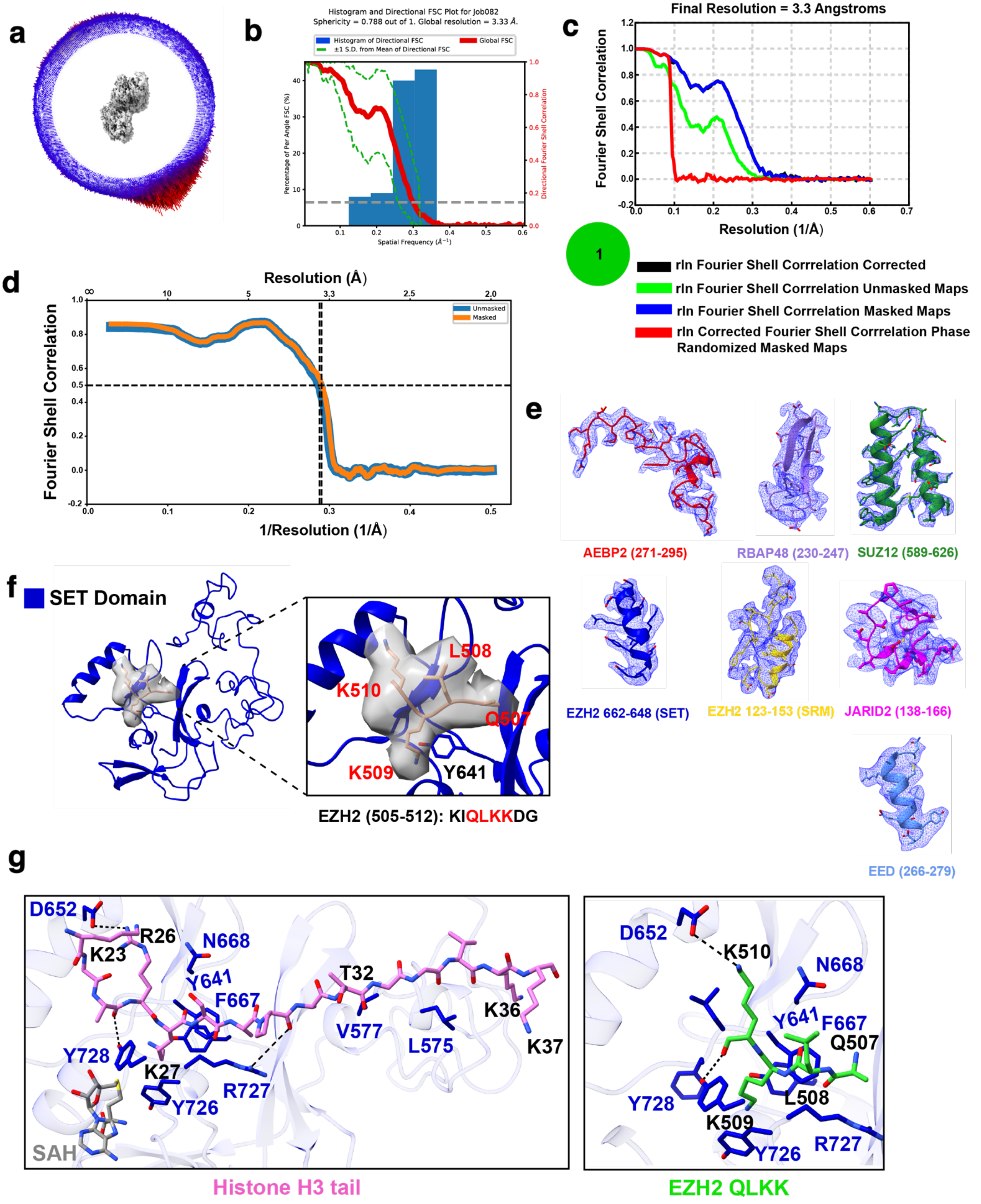


**Supplemental Figure 7. Validation of final EM map and model used in reconstruction. (A)** Angular distribution of the full complex of particles used in final reconstruction. **(B)** 3DFSC for the final map used in reconstruction. **(C)** Gold standard Fourier Shell Correlation for the final refinement in RELION. **(D)** Map-to-model FSC generated with PHENIX. **(E)** Representative high-resolution sections within the final map with the model built in. (**F**) Close-up view of EZH2(SET) domain in the PRC2-AJ119-450 bound to Phe-type activator showing EZH2(SET) domain in the active state with additional density in the catalytic substrate site. EZH2 aa 507-510 region could be modeled into this density consistent with previous reports that this region of EZH2 is a substrate for PRC2 and is automethylated. (**G**) Comparison between Histone H3 tail in the SET domain of EZH2 (6WKR) and the EZH2 QLKK (9ODA) automethylated segment and the associated side chain interactions. SAH was identified in 6WKR but not in 9ODA.

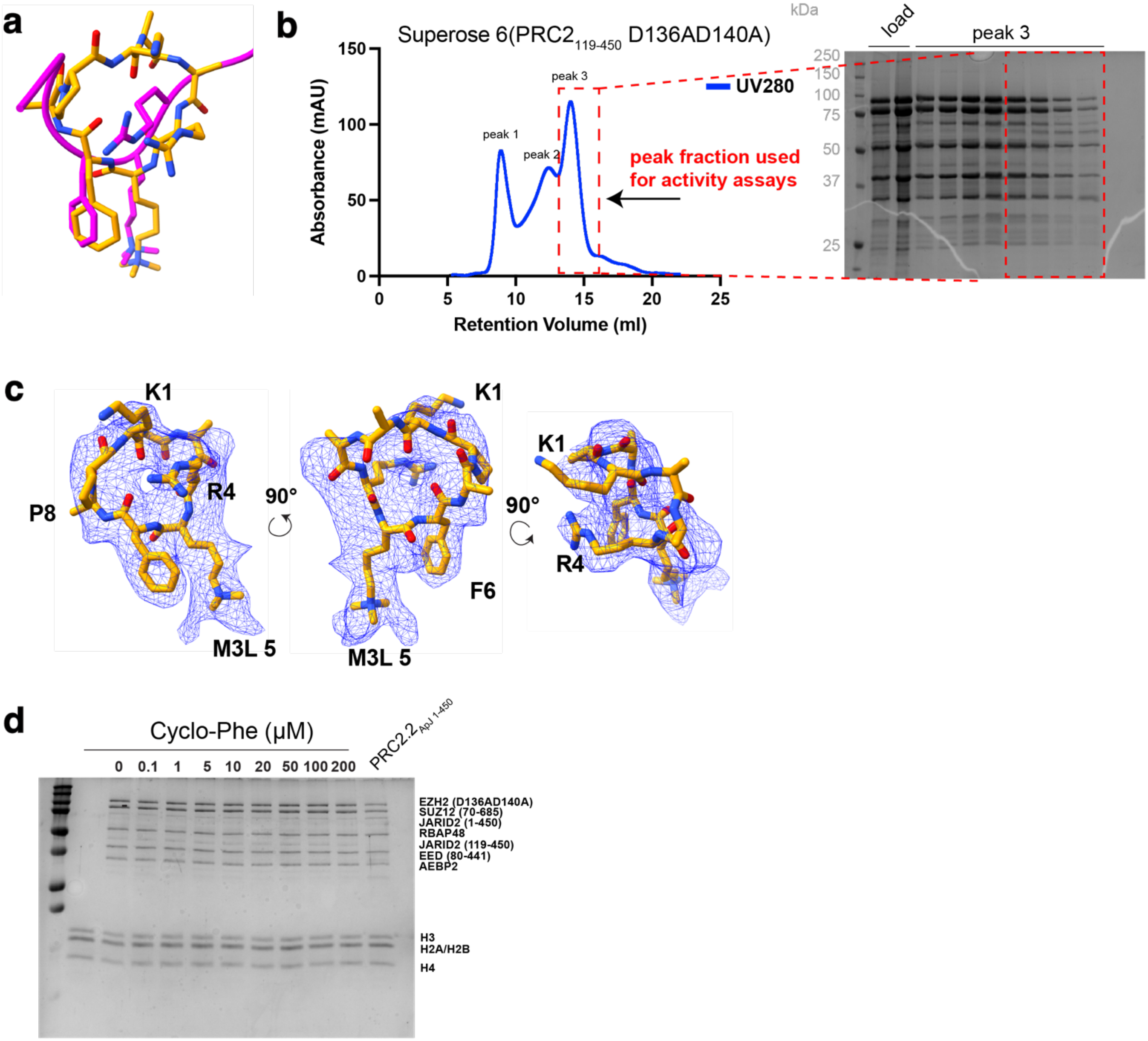


**Supplemental Figure 8. Supplemental data and validation for the cyclic peptide model. (A)** Comparison of Phe-type cyclic peptide model and JARID2K116me3 (aa 114-118) from PDB:6WKR. (B) SDS-PAGE Coomassie stain of purified PRC2$_{119\text{-}450}$ WT and D136A/D140A. Peak 1 is void, peak 2 is dimeric PRC2 observed sometimes due to high-concentrations achieved during sample injection into size-exclusion chromatography. Peak 3 is monomeric PRC2. **(C)** Different orientations of cyclopeptide sitting within Coulomb potential map. (**D**) Representative SDS-PAGE gel Coomassie-stained after Western blot transfer to membrane used in methyltransferase assay of PRC2-AJ119-450 (D136A/D140A).

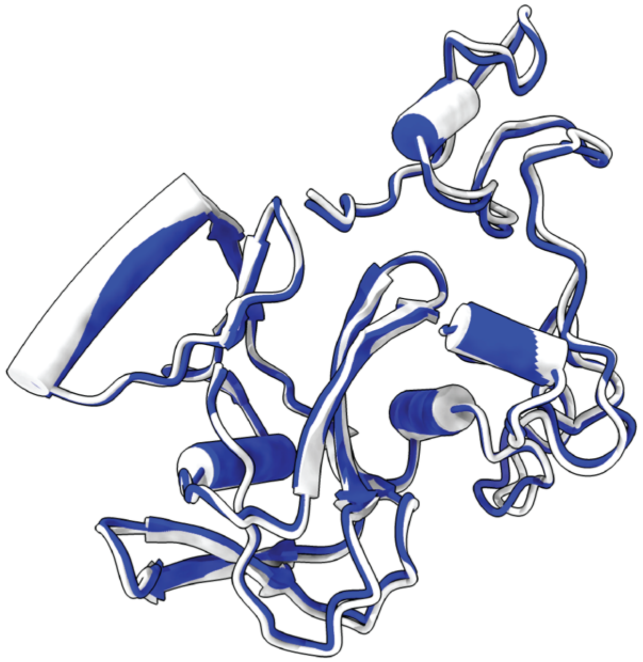

**Supplemental Figure 9. Comparison of the SET domain of 6WKR with the model generated in this study.** Overlay of SET domain shows no noticeable differences (r.m.s.d = 0.78 Å) between the intrinsically allosterically activated PRC2 (PRC2-AJ$_{1-450}$ ; Gray) and PRC2-AJ$_{119-450}$ SET domain (blue) when bound by the cyclic peptide activator.

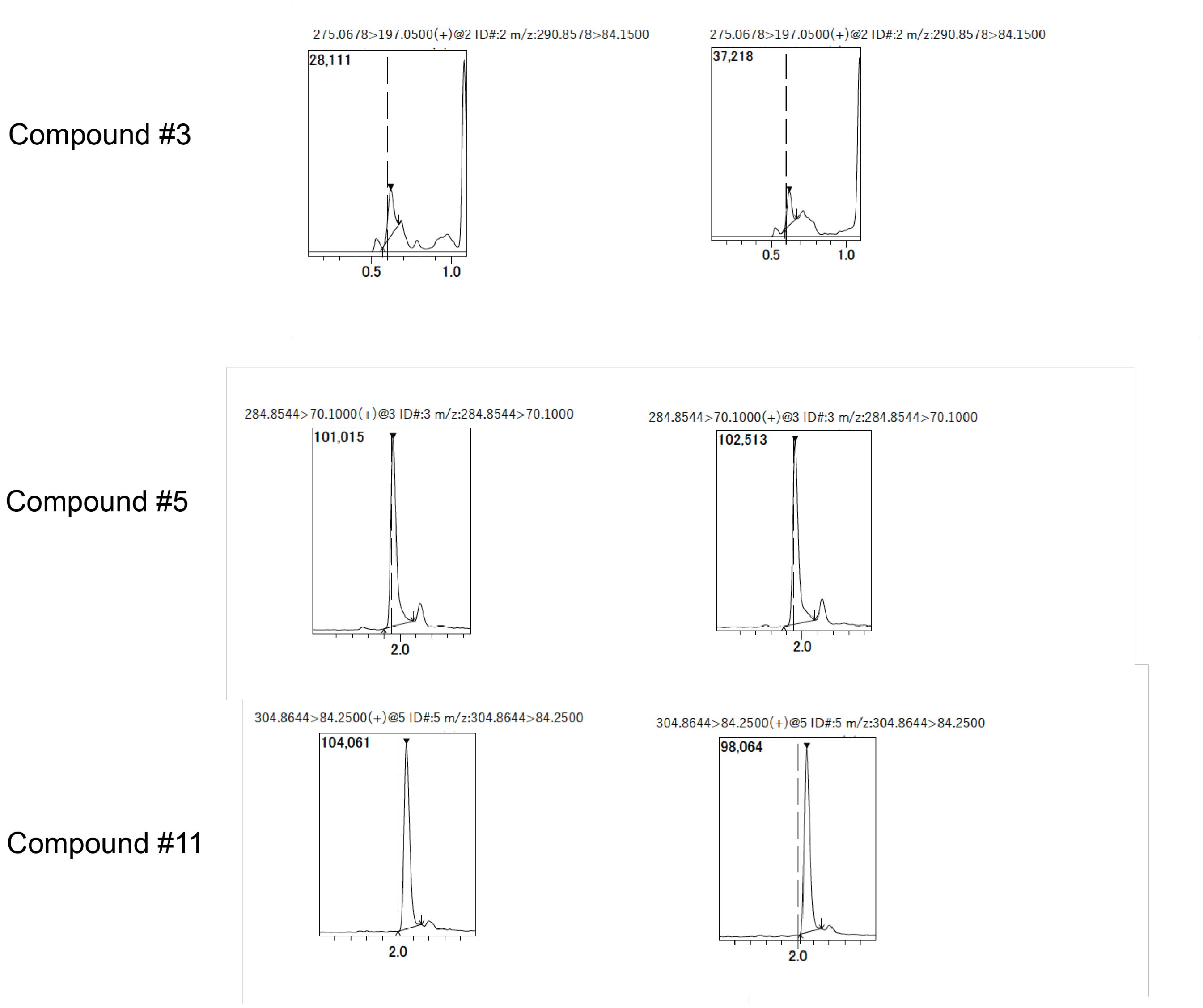


**Supplementary Figure 10.** Mass spectrometry traces for detection of peptide compounds used for mouse plasma stability test at 0 min after incubation.

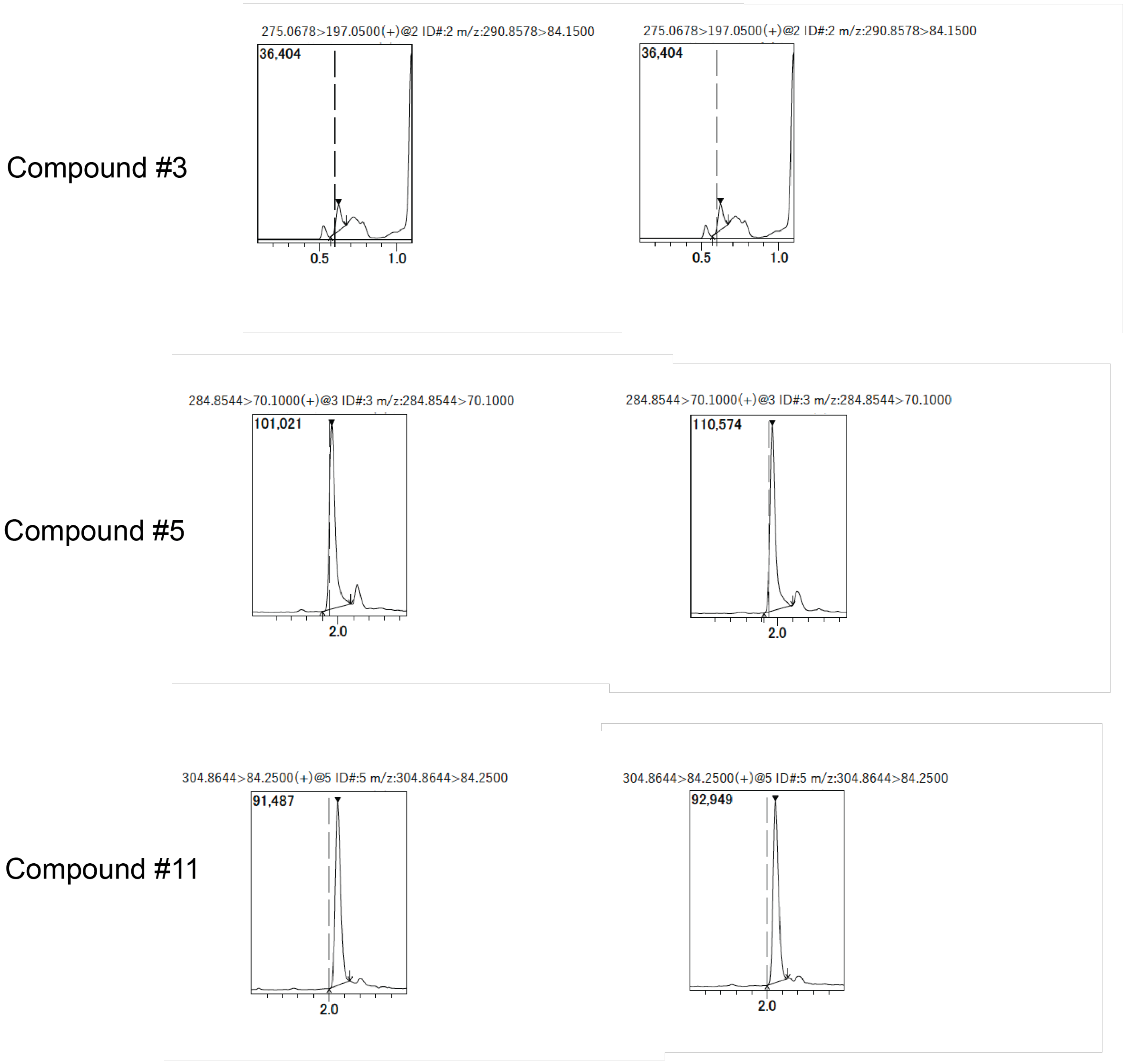


**Supplementary Figure 11.** Mass spectrometry traces for detection of peptide compounds used for mouse plasma stability test at 60 min after incubation.

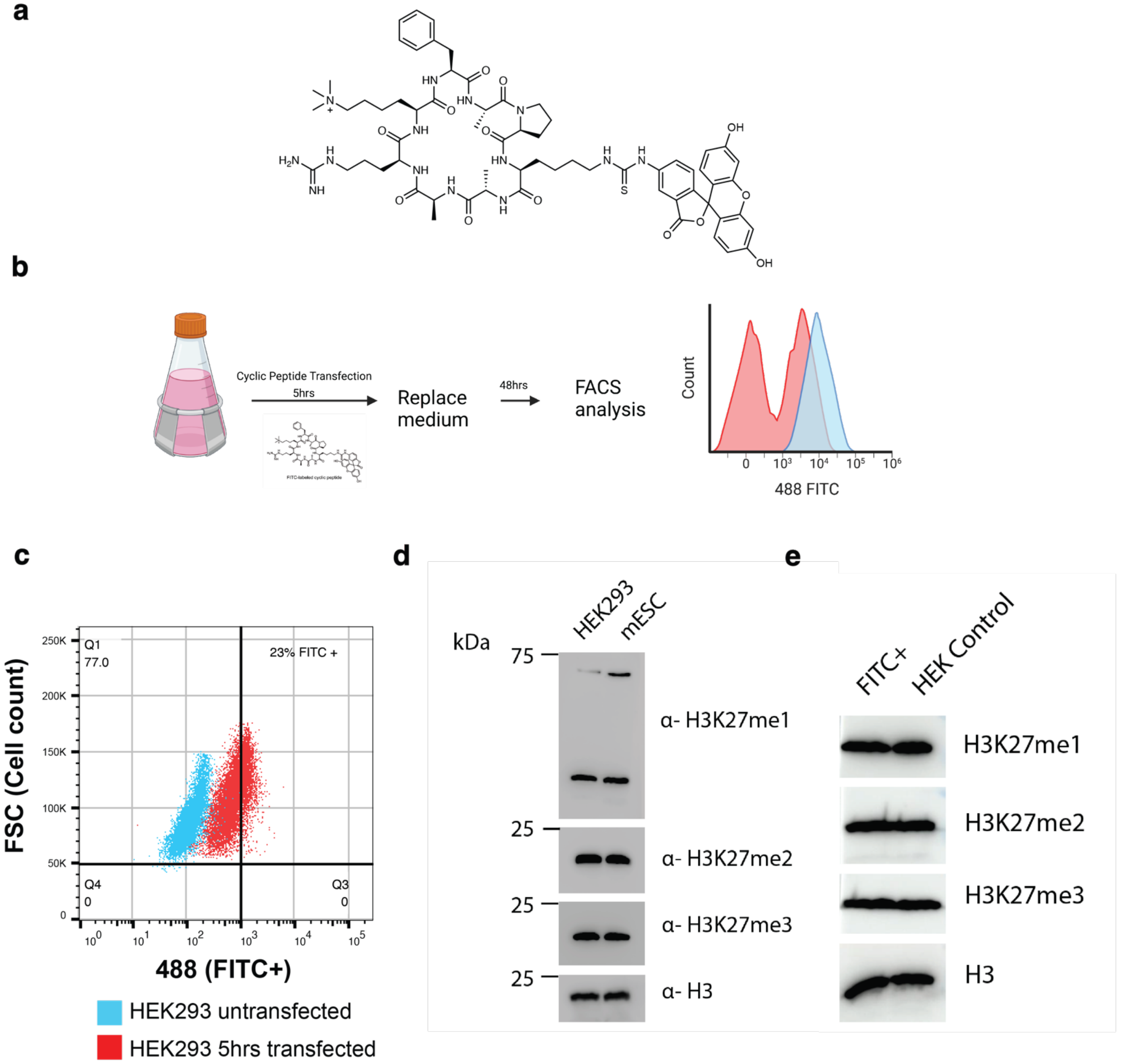


**Supplementary Figure 12.** (a) Structure of the FITC-labeled Phe-type cyclic peptide activator. (b) Schematic of the FACS experiment for isolating FITC+ HEK293 cells. (c) Overlaid cytogram profiles for both HEK293 untransfected DMSO control (blue) and 5 hr transfected (red). (d) Comparison of the H3K27me1/2/3 levels in HEK293 cells versus mESC cells show high levels of these modifications. (e) Comparison of the FITC+ HEK293 cells from FACS sorting (upper right quadrant from (c) – 23% of cells) versus HEK293 control cells show no change in H3K27me1/2/3 levels as control cells already show high levels of these modifications and therefore mask any potential effect of the cyclic peptide activator.

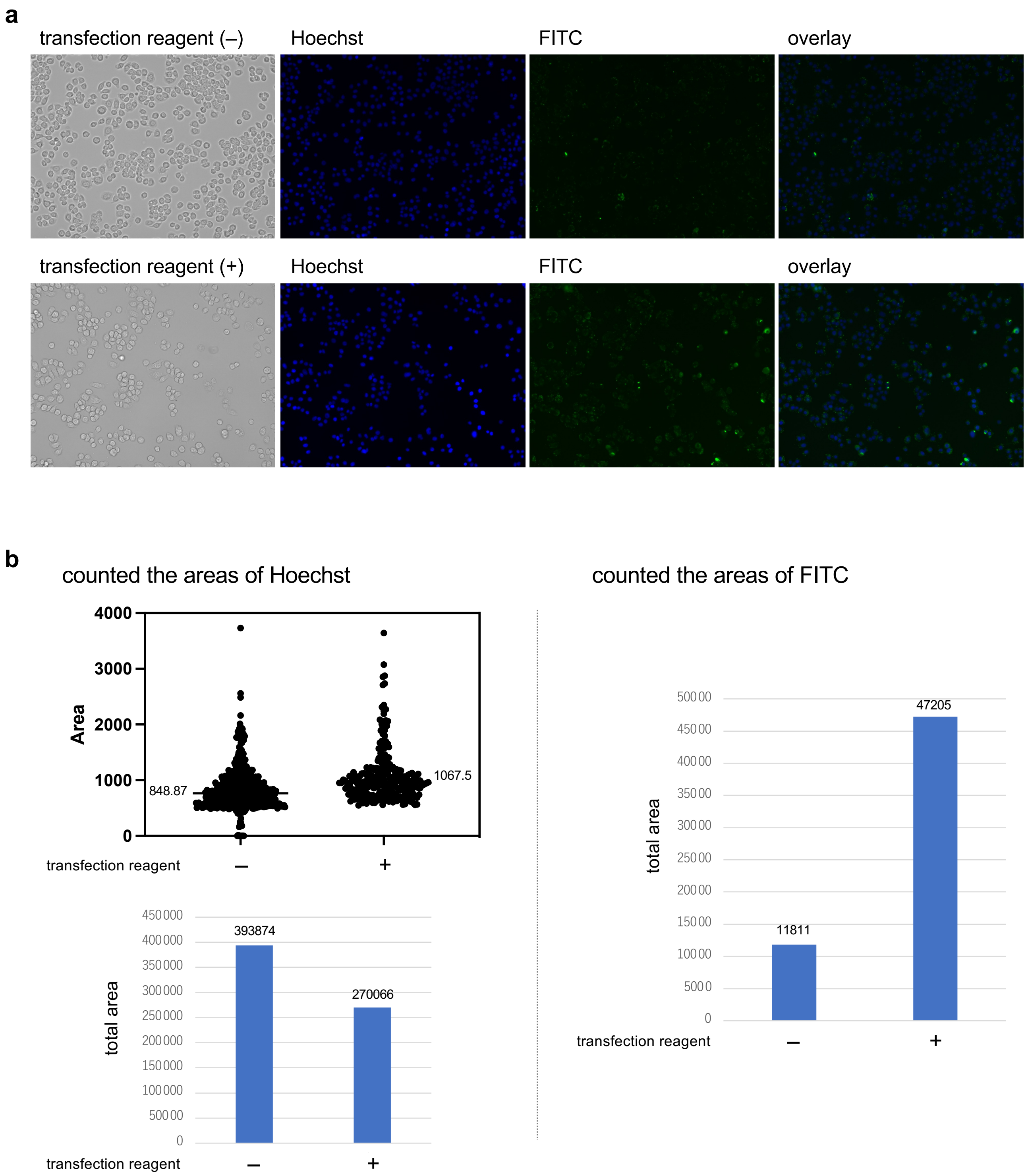


**Supplemental Figure 13. Cellular uptake of FITC-labeled Phe-cyclic peptide.**

(A) Cellular uptake of FITC-labeled Phe-cyclic peptide with HT-29 cells incubated for 1 h, with or without transfection reagents. (B) The areas of Hoechst (blue) and FITC (green) were counted. The area of Hoechst of each of the cells and the averages are shown (left). The total areas of Hoechst (left) and FITC (right) are depicted. The images were counted by ImageJ 1.54g.

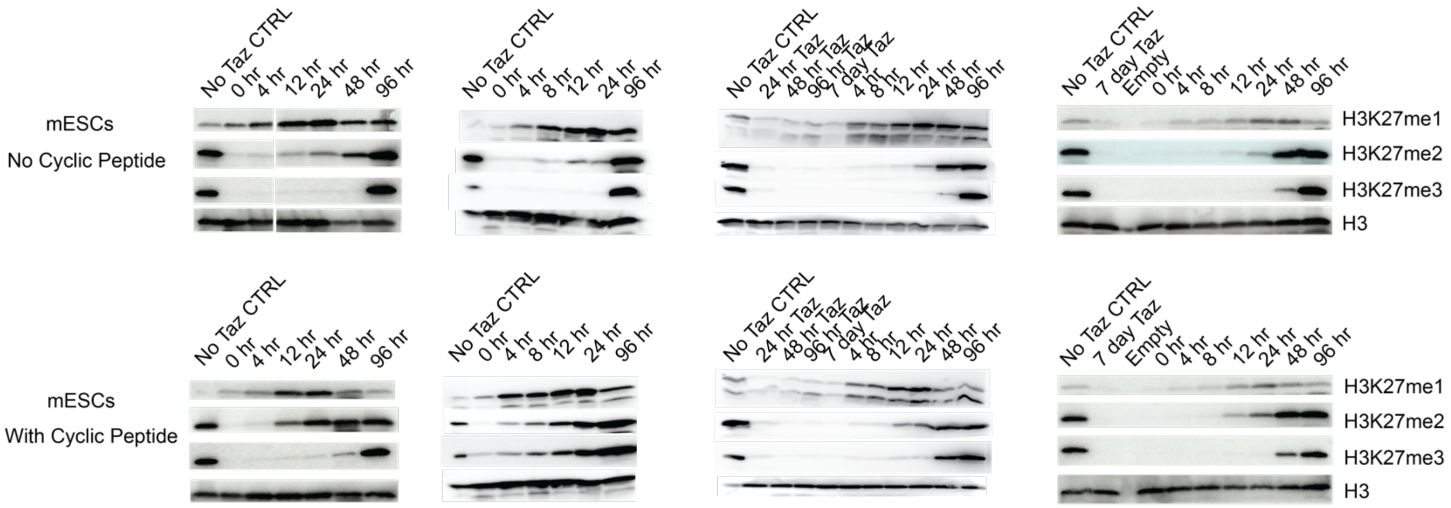


**Supplementary Figure 14.** Western blot assays of H3K27me1/2/3 in mESCs after Tazemostat treatment and recovery either with or without cyclic peptide activator.

### Supplemental Figure Raw Data (gels)

#### Fig. 2C

##### H3 trimethylation assay

Ser-type H3K27me3 uncropped immunoblot

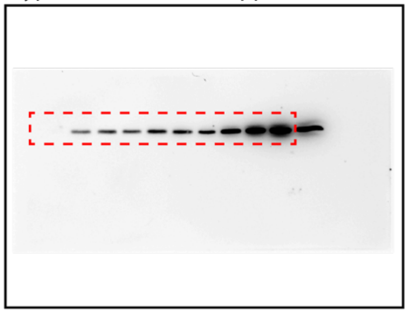

Phe-type H3K27me3 uncropped immunoblot

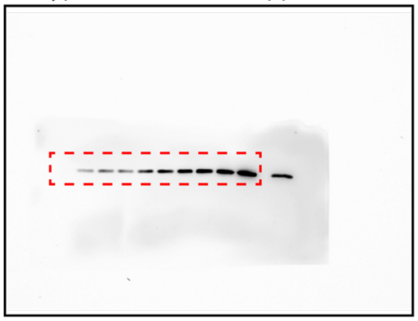

Histone H3 uncropped immunoblot

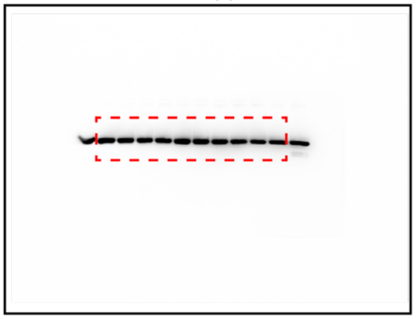

#### Fig. 2D

##### Nucleosome trimethylation assay

Ser-type H3K27me3 uncropped immunoblot

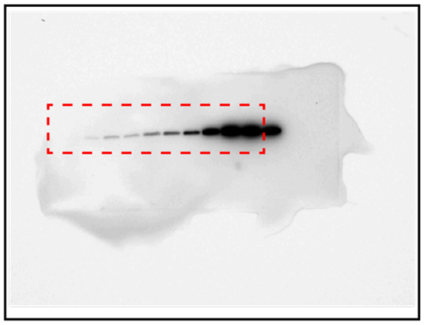

Phe-type H3K27me3 uncropped immunoblot

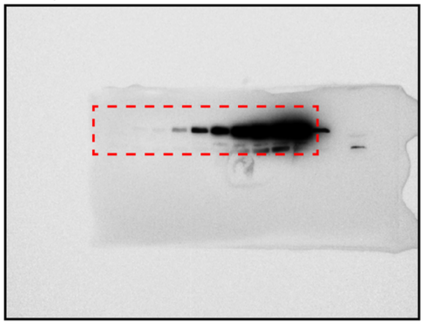

Histone H4 uncropped immunoblot

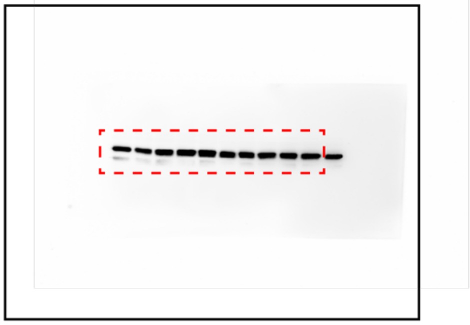

#### Fig. 3C

##### Nucleosome trimethylation assay

Phe-type H3K27me3 uncropped immunoblot

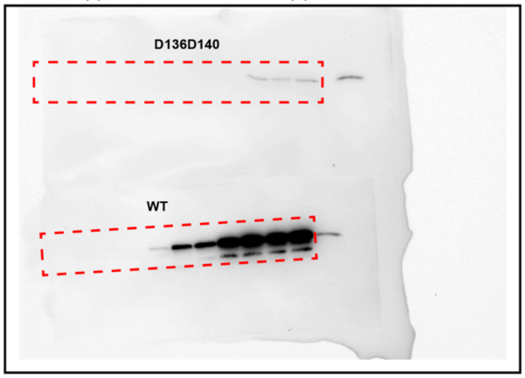


Histone H4 uncropped immunoblot

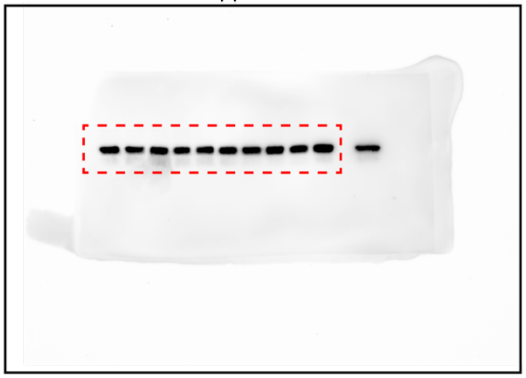

**Fig. 4A**

H3K27me1 uncropped immunoblot

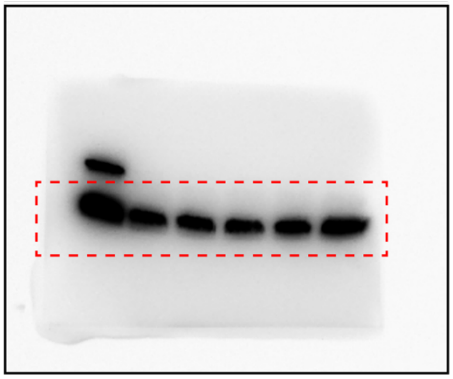

H3K27me2 uncropped immunoblot

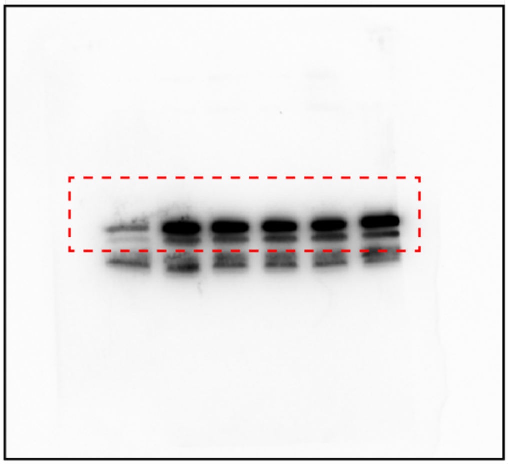

H3K27me3 uncropped immunoblot

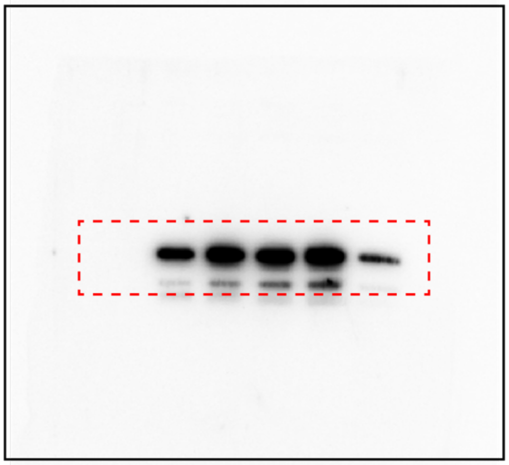

Histone H4 uncropped immunoblot

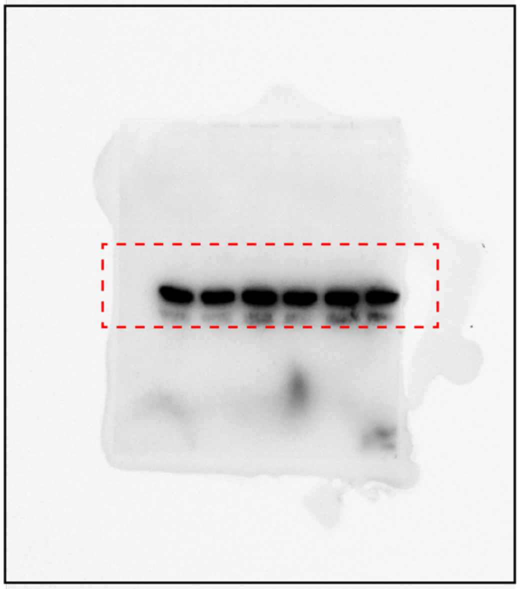

**Extended Table 1.** Summary of cryo-EM structure determination of PRC2-AJ119-450 bound to Phe-type cyclic peptide activator

| **Samples** | **PRC2-AJ$_{119-450}$ – Cyclo Phe** |
|---|---|
| **Data collection and processing** | |
| Microscope | Titan Krios 3 |
| Camera | Gatan energy filtered K3 |
| Energy filter | BioContinuum |
| Magnification | 105 kx |
| Voltage (keV) | 300 |
| Electron exposure ($e^-$/Å) | 50 |
| Defocus range (μm) | -0.8 to -2.5 |
| Pixel size (Å) | 0.825 Å/pix (0.4125 Å/pix for super res.) |
| Symmetry imposed | C1 |
| Initial particle images (no.) | 4,939,638 |
| Final particle images (no.) | 72,000 |
| Map Resolution (Å) | 3.3 (local)<br>3.7 (global) |
| FSC threshold | 0.143 |
| **Refinement** | |
| Initial model used (PDB code) | 6WKR |
| Model resolution range (Å) | 3.3-8 Å |
| Map sharpening B factor (Å$^2$) | -55.8 |
| Model composition | |
| Nonhydrogen atoms | 25494 |
| Protein residues | 1846 |
| Nucleotides | 0 |
| B factors (Å$^2$) | |
| Protein | |
| Nucleotides | |
| R.m.s. deviations | |
| Bond lengths (Å) | 0.002 (0) |
| Bond angles (°) | 0.473 (1) |
| **Validation** | |
| MolProbity score | 1.63 |
| Clashscore | 4.04 |
| Poor rotamers (%) | 1.27 |
| Ramachandran plot | |
| Favored (%) | 94.84 |
| Allowed (%) | 5.11 |
| Disallowed (%) | 0.06 |
